# Comparative Biology at Single-Cell Resolution: Rigorous Matching of Atlases for Cross-Species Analysis

**DOI:** 10.64898/2026.02.18.706600

**Authors:** Marc-Antoine Jacques, Berthold Göttgens, John C. Marioni

## Abstract

Single-cell transcriptomics has revolutionised developmental biology by providing an unprecedented, fine-grained view of cellular lineages. However, our ability to compare species and distinguish universal from species-specific developmental principles remains limited by biological and technical variability. To address this, we introduce RIMA (RIgorous Matching of Atlases), a method for quantitatively comparing transcriptomic atlases across species at near-single-cell resolution. RIMA uses a novel computational approach to identify matching cell states across atlases and leverages this to enable quantitative comparative analyses.

Applied to gastrulation in mouse, rabbit, and macaque, RIMA recapitulates a developmental hourglass pattern, identifying a molecular similarity bottleneck at the onset of organogenesis. It further uncovers conserved developmental programmes, including a core set of transcription factors driving epithelial-mesenchymal transition, and highlights transcriptional boosts of erythroid differentiation genes that are conserved across species but exhibit shifted onset timing. Furthermore, RIMA enables cross-species prediction of gene expression, augmenting sparse atlases and correcting differences between model and target organisms.

Beyond cross-species comparisons, RIMA’s framework extends naturally to any setting where large systematic differences exist across datasets, including *in vitro* to *in vivo* comparisons, opening new avenues for improving biological models and advancing translational research.

## Introduction

Developmental biology has long relied on a range of model organisms to distil the broad principles governing embryonic development. The use of multiple species has facilitated the distinction between general and species-specific processes, as well as the utilisation of the experimental advantages offered by different systems^1–4^.

Single-cell sequencing has revolutionised developmental biology by enabling the construction of comprehensive gene expression atlases in multiple species, addressing long-standing questions about progenitor sequences and differentiation trajectories within individual organisms^4–10^. However, comparing these atlases to identify common mechanisms remains challenging due to their complexity, spanning different species, non-overlapping developmental stages, complex cell type composition, and divergent genomes of variable quality. Thus, most cross-species analyses have relied on simple approaches, such as matching the annotations of entire cell clusters or comparing predefined marker genes. Although insightful, this sacrifices the potential gains offered by single-cell resolution by averaging cells or focusing on known gene subsets. Other attempts relied on single-cell integration methods, which aim to align shared cell states by removing technical and sample-specific (including species-specific) variability. However, these approaches were mostly developed for batch-effect removal and they often struggle with large biological shifts, such as the presence of multiple species^11–13^. More specialised methods have been developed^14,15^, but often produce features that are difficult to interpret and are highly sensitive to the evolutionary distance between species^16^. Alternatively, perturbation prediction models^17,18^, including foundation models^19,20^, have been proposed to analyse cross-species transitions, but recent evidence questions the accuracy of their prediction^21^.

Here we developed an alternative approach, Rigorous Matching of Atlases (RIMA), to compare single-cell transcriptomics atlases, including cross-species comparisons, with minimal assumptions and interpretable outputs. Critically, our pipeline does not compromise single-cell resolution and does not rely on the construction of integrated embeddings. Instead, it builds an explicit matching between atlases at the level of cell neighbourhoods, based on their transcript counts. The sole assumption of RIMA is that a meaningful distance between neighbourhoods can be computed to capture equivalent cellular identities across atlases. RIMA uses a novel procedure to identify candidate matches before returning a one-to-one matching of neighbourhoods across atlases. This procedure is scalable, robust to different cell type compositions and does not rely on manual annotation. The results can be mined at multiple scales, from whole atlases to individual genes and cell clusters, and reveal quantitative differences between atlases. In the context of developmental biology, RIMA is particularly well-suited for studying divergence and conservation, but it can be extended to other contexts, especially those where integration is difficult.

In this work, we introduce the RIMA methodology through a case study comparing gastrulation across mammals (mouse^22^, rabbit^6^ and macaque^7^). We first demonstrate RIMA’s robustness by comparing the mouse and rabbit atlases, showing that it recapitulates known proximities between cell types and developmental stages. We then use this explicit matching at a finer resolution to compare trajectories of erythrocyte development and reveal functional as well as temporal conservation between the two species. Next, we incorporate a macaque atlas and identify a core set of 83 transcription factors, mostly associated with epithelial-mesenchymal transition, whose activity is strongly preserved in matching cell states. Finally, we show that RIMA’s matching can be used to predict transcript counts across species, correcting known model limitations and augmenting sparse atlases from less-characterised species by leveraging extensive datasets from better-characterised organisms.

## Results

### RIMA: a versatile pipeline for matching and quantitatively comparing atlases at near-single-cell resolution

RIMA performs atlas matching at the scale of cell neighbourhoods^23,24^ thus retaining single-cell resolution, mitigating sequencing noise and improving scalability for larger atlases. RIMA begins by calculating all pairwise neighbourhood similarities across species based on the neighborhoods’ mean gene expression (Fig. 1). All pairwise similarities can be represented as a weighted bipartite graph, where each pair of neighbourhoods (nodes) across species (sets) is connected by a weighted edge. Throughout this study, each neighborhoods’ gene expression is defined as the average gene expression of the cells within that neighbourhood. We use Spearman correlation as the similarity measure, which was reported as a good distance to capture cell types^25^, and which we found returns robust results for our matching across species. This is likely because the ranking of gene expression is more important to define cellular identity than absolute transcript levels, which are less conserved across species and more susceptible to technical variation.

**Figure 1:**
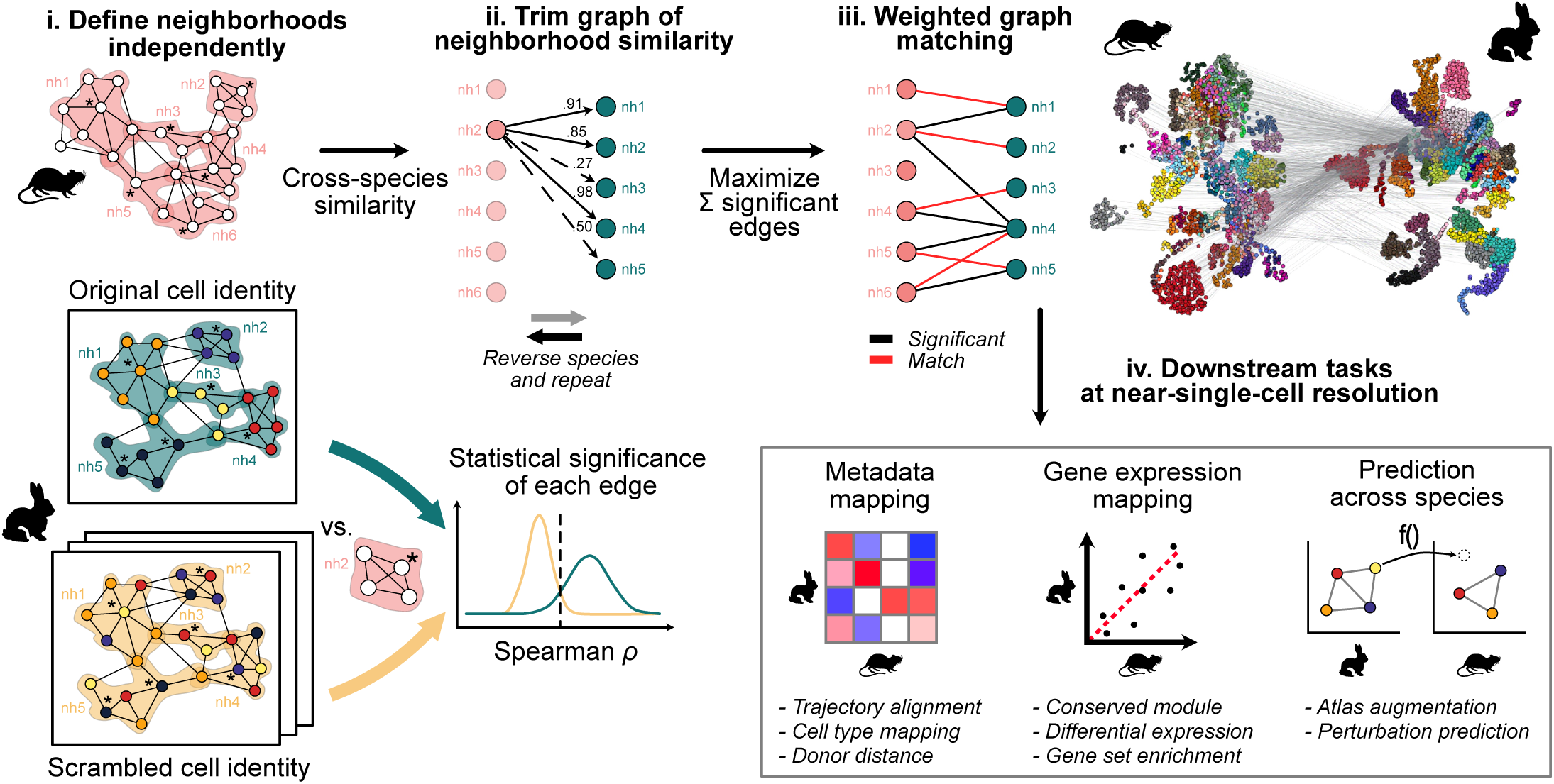
Overview of RIMA. RIMA matches neighbourhoods across single-cell transcriptomics atlases by first assessing the statistical significance of the similarity between every pair of neighbourhoods across the two atlases. For each edge linking two neighbourhoods, its significance is estimated by randomly resampling cells from one atlas and comparing the similarity between the original neighbourhoods and the similarity between the original and resampled neighbourhoods. This process is repeated by resampling the second atlas, yielding a second p-value for each edge. The two P-values are combined to determine the statistical significance of each edge. The graph comprising only statistically significant edges is resolved using weighted graph matching, which pairs neighbourhoods across atlases for use in downstream analysis. In the right panel, an example matching between the mouse and the rabbit gastrulation atlases is presented (see Fig. S1), where only half of the neighbourhoods matches are shown for clarity.

RIMA then prunes insignificant edges from the bipartite graph to retain only candidate neighbourhood matches by comparing the edges’ similarity to a “null population”. Specifically, edge significance is determined by comparing the distribution of similarities between original neighbourhoods and those with scrambled neighbourhoods, where cell identities have been randomly resampled. This enables the calculation of an empirical, directional p-value for each edge. To ensure that the edge is also significant against a “null population” formed by scrambling the other species, the procedure is repeated in the opposite direction, and the two p-values of each edge are combined using Simes’ method^26^. Edges with a combined p-value below a specified significance threshold are considered candidate matches and passed to the next step. Optionally, the resampling of neighbourhoods can be weighted to account for imbalances in neighbourhood metadata (e.g., a very large batch or a rare cell type), thereby generating truly unspecific, scrambled neighbourhoods. This filtering step ensures that only relevant neighbourhood pairs are passed on to the matching step (Fig. 2a,b).

**Figure 2:**
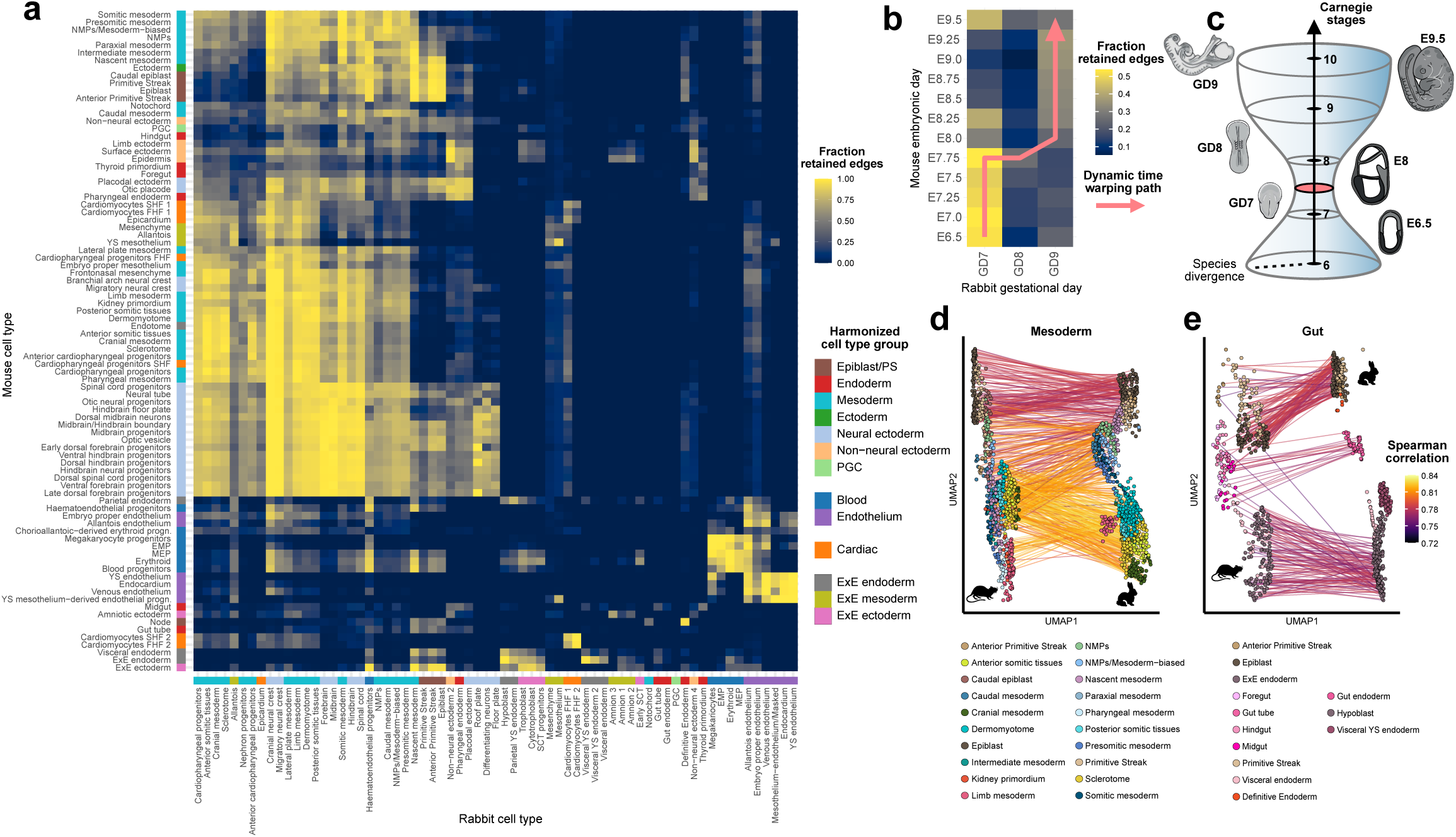
Whole-embryo comparison of mouse and rabbit gastrulation with RIMA. **a-b**, Fraction of neighbourhood-neighbourhood edges retained by RIMA. Rows and columns represent the cell type, (**a**), or developmental time, (**b**), annotations of neighbourhoods in both species. Initially, every pair of neighbourhoods is connected between the two species, and statistically insignificant edges are trimmed. The pink arrow indicates the optimal DTW alignment path between species’ stages. **c**, Schematic of the developmental hourglass concept. The identified bottleneck of molecular similarity is highlighted in pink. **d-e**, Pairs of neighbourhood matched by RIMA, subsetted to the mesoderm (**d**) and gut lineage (**e**). Matches are coloured by Spearman correlation between the mouse (left) and rabbit (right) neighbourhoods.

In the final step, RIMA resolves the resulting graph into a one-to-one matching using maximum weight matching, which maximises the sum of the weights of the matched edges. This global approach avoids the “kissing effect” which would be introduced by matching every neighbourhood to its most similar counterpart in the other set without integration (Fig. S2, Supplementary Note 1). The “kissing effect” leads to neighbourhoods of one cluster tending to match those on the surface of the other cluster. This creates “clumps”, where many neighbourhoods from one atlas are mapped to a few in the other, leading to a discrete matching that does not capture the continuity of the biological landscape (Fig. S2c).

RIMA is an unsupervised pipeline, where nearly every step is non-parametric and free from biases in metadata or parameter choice. We demonstrate below that RIMA enables the comparison of atlases in developing embryos from diverse species – a particularly challenging task due to the mixing of different species, the presence of numerous cell types at varying degrees of differentiation, and the inclusion of multiple time-points derived from several sequencing rounds in many individual embryos.

### RIMA’s matching of gastrulation atlases recapitulates cell type proximity and highlights a developmental hourglass similarity profile at the molecular level

Gastrulation is a critical step of animal development during which the three germ layers (endoderm, mesoderm, ectoderm) are established and then rapidly give rise to a multitude of tissues, shaping the future body plan. Many core elements of gastrulation have long been known to be critically conserved across species through anatomical descriptions, and recent single-cell atlases in a few species have also revealed striking molecular similarities^4,6–9^. However, cross-species comparisons at the single-cell level have typically been limited to high level tasks such as verifying cell type consistency through label transfer, and inspecting the expression of known, individual marker genes. Here, we revisit cross-species similarity at the molecular level with RIMA, making full use of the resolution offered by single-cell data.

We applied RIMA to compare two of the most comprehensive publicly available atlases of gastrulation: a rabbit (*Oryctolagus cuniculus*) atlas^6^ and a mouse (*Mus musculus*) atlas^22^ (Fig. S1). Both atlases span from the onset of gastrulation to early organogenesis, were sequenced using the same technology (10X Genomics Chromium), and were processed with consistent pipelines.

We found that RIMA was able to map every neighbourhood from the mouse (n=3448) to a neighbourhood in the rabbit atlas (n=5254), highlighting the conservation of gastrulation across these two species. By inspecting the map of trimmed neighbourhood-neighbourhood edges (Fig. 2a,b), we observed that a large fraction of edges were deemed insignificant, with combined p-values above 0.05 after adjustment for family-wise error rate. From the perspective of cell types, the trimmed map captures known cell type lineages and highlights specific areas, such as the haematoendothelial lineage, and cardiomyocytes, which are highly separated from others. This map also suggests a higher proximity between early tissues (i.e. epiblast, primitive streak) and ectoderm-derived cells than with the other germ layers. This observation could be related to the fact that ectodermal cells are derived from epiblast cells which – unlike mesoderm and endoderm – do not undergo epithelial-to-mesenchymal transition (EMT). We also analyzed the trimmed map between embryonic stages and identified a bottleneck connecting gestational day 8 (GD8) in rabbit to embryonic day 7.75 (E7.75) in mouse (Fig. 2b). The timing of this bottleneck is consistent with previous observations^4^ describing an evolutionary hourglass model (Fig. 2c), which states that species go through a transient stage of high molecular similarity during early development. Our analysis suggests that this stage arises at the onset of organogenesis, immediately prior to the commitment of cells to downstream lineages (Fig. S4).

We then inspected the matching of known developmental trajectories (Fig. 2d,e, Fig. S3). We found that the mesodermal lineage was particularly well-conserved across the two species, as highlighted by the strong agreement of matched cell types and very high similarity values. The gut lineage, however, displayed greater variability. Although most cell types had a corresponding match in the other species, the rabbit visceral endoderm lacked a counterpart, and a few mouse gut neighbourhoods were mapped to the rabbit yolk sac and definitive endoderm. In the mouse, the gut tube forms through the intercalation of cells from the visceral endoderm (yolk sac-derived) and the embryo-proper endoderm. Hence, RIMA’s mapping could reflect that while the mouse gut cells retain a signature from both origins^27^, our results would be consistent with such direct intercalation not happening in the rabbit, or a failure to retain distinct signatures.

### RIMA’s matching reveals cross-species conservation of erythrocyte lineages development and subtle divergence of gene expression dynamics

#### The conservation of erythrocyte formation is driven by expression programs linked to general cell stemness and haematopoietic metabolism

Embryonic haematopoiesis emerges through a tightly controlled spatio-temporal process during development. The first haematoendothelial progenitors are formed in two successive waves in the yolk sac around E7.25 and E8.25 in the mouse. The increasingly differentiated progenitors then migrate to the liver where they produce definitive erythrocytes and myeloid cells around E10.5, until the bone marrow production becomes functional^28^.

Having established that RIMA produces a biologically-relevant matching between mouse and rabbit gastrulation, we next sought to provide a fine-grained comparison of the erythrocyte differentiation trajectory between the two species and unravel similarities and differences in the underlying molecular regulatory landscape. We first observed that the neighbourhoods associated with these trajectories were matched with very high similarity values (Fig. 3a), and that pseudotimes derived independently from both species were nearly linearly associated across mapped neighbourhoods (Fig. 3b, S2). Together, these results suggest that the establishment of haematopoiesis is extremely well-conserved from epiblast to differentiated erythrocytes, consistent with previous studies^6,29^.

**Figure 3:**
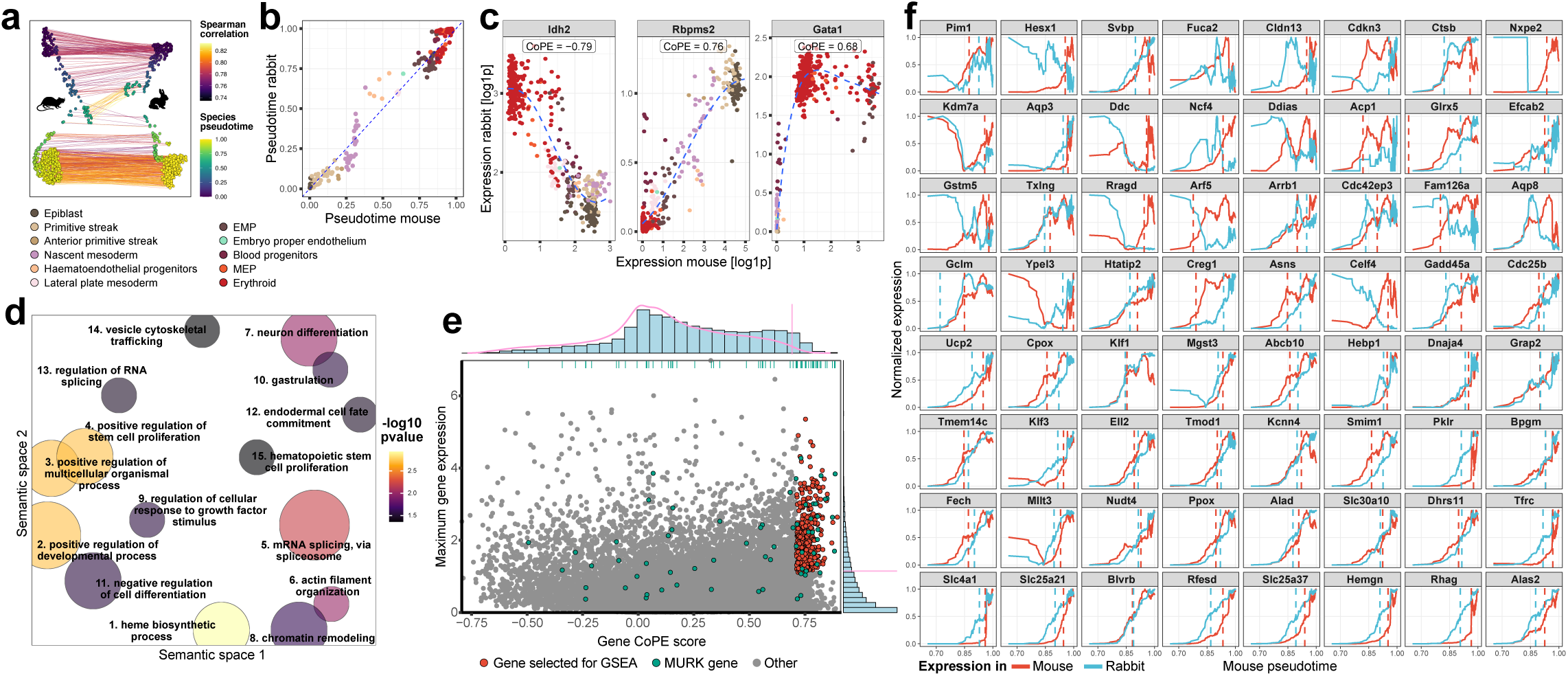
Comparison of erythrocyte development between mouse and rabbit. **a**, RIMA matching of mouse and rabbit erythrocyte lineage neighbourhoods. Edges coloured according to Spearman correlation of the edge. **b**, Erythrocyte lineage pseudotimes of the matched pairs from **a**, colored by mouse cell type annotation. **c**, Representative examples of genes with high and low CoPE scores, as well as the haematopoietic marker Gata1. **d**, GSEA of genes selected in **e** with GO biological processes. The semantic space gathers similar terms under 15 representative terms for visualisation. Size of the dots indicates the number of terms in the cluster, colored by enrichment significance. **e**, Selection of highly conserved and expressed genes for GSEA. The vertical axis indicates the maximum raw count observed in any erythrocyte neighbourhood, either in rabbit or mouse, whichever is higher. Marginal distributions indicated as histograms, pink lines indicate selection cutoffs for GSEA. The CoPE score cutoff was determined by randomly permuting gene identities ten times, calculating CoPE scores using the permuted genes, and selecting the value below which 99% of the permuted CoPE scores fell (see Methods). The expression cutoff was manually set to 2 counts. Green rugs and dots indicate MURK genes’ position. **f**, MURK gene expression along mouse erythrocyte pseudotime. Rabbit neighbourhoods were projected onto the mouse pseudotime using the pseudotime of their paired mouse neighbourhood. Vertical dashed lines indicate the midpoint of successful sigmoid fits. Gene expression levels were normalised between 0 and 1, and smoothed with rolling mean.

We then sought to characterize the gene expression programmes that were most conserved between the two species. For this, we defined a Correlation of Paired Expressions (CoPE) score for each gene. This score is defined as the Spearman correlation of the expression levels across the matched neighbourhoods. Our rationale for this metric is that genes whose expressions are concordant between matched cellular identities are likely to be important differentiation markers and more conserved^30^. As expected for orthologs, we observed a right-skewed distribution for this metric (Fig. 3e), indicating that, across neighbourhoods, expression levels for most genes varied consistently between the two species. Many genes had CoPE scores around 0 or below 0.25, suggesting that only a small subset of genes were involved in the progression towards mature erythrocytes. More surprisingly, we also observed that some genes were heavily negatively correlated, hinting at cross-species divergence. Upon manual inspection of individual genes (Fig. 3c), we found that the gene with the lowest CoPE score (−0.79), Idh2, is mitochondrial. Hence, its expression could be expected to vary more freely between species due to the semi-autonomy of the organelle, and the higher mutation rate of mitochondrial genomes^31^. On the other hand, high CoPE scores captured genes that are representative of genes that are both upregulated and downregulated along the trajectory, for example Gata1 (CoPE = 0.68; 92th centile), a classic marker of erythroids, and Rbpms2 (CoPE = 0.76; 98th centile), an inhibitor of BMP signaling, a major pathway promoting haematopoiesis.

We then characterized the most conserved genes by selecting those with high CoPE scores and high expression levels (Fig. 3e), and ran a gene set enrichment analysis (GSEA) using the GO biological processes sets^32,33^ (Fig. 3d, Supplementary Table S1). We found that the most conserved genes along this trajectory were strongly associated with terms related to stem cells and erythrocyte metabolism (e.g. heme metabolism). In contrast, the negatively correlated genes were only enriched in generic housekeeping terms, such as transcription regulation and energy metabolism, including mitochondrial genes. This disparity likely reflects the dominance of cell-type-specific signals over baseline cellular functions in driving the beginnings of haematopoiesis and the resulting mapping of RIMA.

#### Paired neighbourhoods allow for the alignment of development trajectories and reveal conserved but shifted transcriptional boosts

Developmental trajectories are characterized by gradual or sudden changes in expression of marker genes. A group of important regulators of erythrocyte differentiation in the mouse, called multiple rate kinetics (MURK) genes, was identified on the basis of their sharp increase in expression during differentiation^34^. To date, this peculiar dynamic has not been described in other organisms. Using RIMA’s matching scheme, we sought to align the erythrocyte developmental trajectories across both species and investigate whether such boosting was also present, and synchronized, in the rabbit.

We used RIMA’s matching scheme to establish a common coordinate system that reflects the progression towards erythrocyte states in both species. To obtain these coordinates, we first isolated the erythrocyte lineage cells in both species and ordered them along species-specific pseudotimes. We then assigned a pseudotime to each neighbourhood by averaging the pseudotimes of the cells that composed them. We then looked up RIMA’s mapping to get a direct correspondence between species-specific pseudotimes. This allowed us to describe progression in the rabbit trajectory using mouse pseudotime units, and vice versa (Fig. 3f).

We first noted a nearly linear correspondence between the species-specific pseudotimes (Fig. 3b), hinting at a strong conservation of the sequence and intermediates leading to erythrocyte differentiation. We then constructed expression time-series for both species, using the mouse pseudotime as the reference “temporal” axis. Finally, we identified the boosting genes by fitting a logistic function to the last segment of these time-series, where cells are committed to the erythroid-myeloid lineage and most MURK genes are expressed. Out of the 64 original MURK genes with a one-to-one ortholog between the species, we confirmed a sigmoidal boost of transcription in 60 genes in the mouse and found that 49 MURK orthologs also exhibited boosting in the rabbit (Fig. 3f). Although the boosting dynamics of MURK genes appeared preserved, we noted that their onset (as quantified by the sigmoid midpoints) was often shifted between the two species (Fig. 3f). This suggests that, despite being a highly conserved process, the establishment of embryonic haematopoiesis might be established through similar, yet functionally distinct intermediary steps in each species. Additionally, a few genes showed inverted expression profiles, with a decrease of expression along trajectories (e.g. Celf4, Gstm5) in the rabbit. This may indicate that these genes are under the control of transcriptional programs involved in erythrocyte development in the mouse, but that they are not directly involved in the process.

### Mammalian gastrulation is driven by a small core of conserved transcription factors

Next, we sought to expand our comparison of gastrulation to other mammals by adding an atlas corresponding to early development of the crab-eating macaque (*Macaca fascicularis*)^7^, a primate that has diverged from rodents and lagomorphs about 70 million years ago^40^ and is one of the closest species to human that has been comprehensively sequenced. Using the three species, we investigated which gastrulation processes were most conserved by analysing the expression of the transcription factor (TF) regulons contained within the mouse Collectri gene regulatory network compendium^41^. To do this, we applied RIMA to all pairs of species and extended the CoPE score from individual genes to regulons by averaging the CoPE scores of all target genes regulated by a given TF (Fig. 4a). Each regulon’s CoPE score was then compared to random samples of genes of equivalent size to determine whether the regulon was significantly more conserved or divergent than expected (Fig. 4b,c). Notably, the random samples are also formed by orthologs common to the three species, and hence constitute a conservative background.

**Figure 4:**
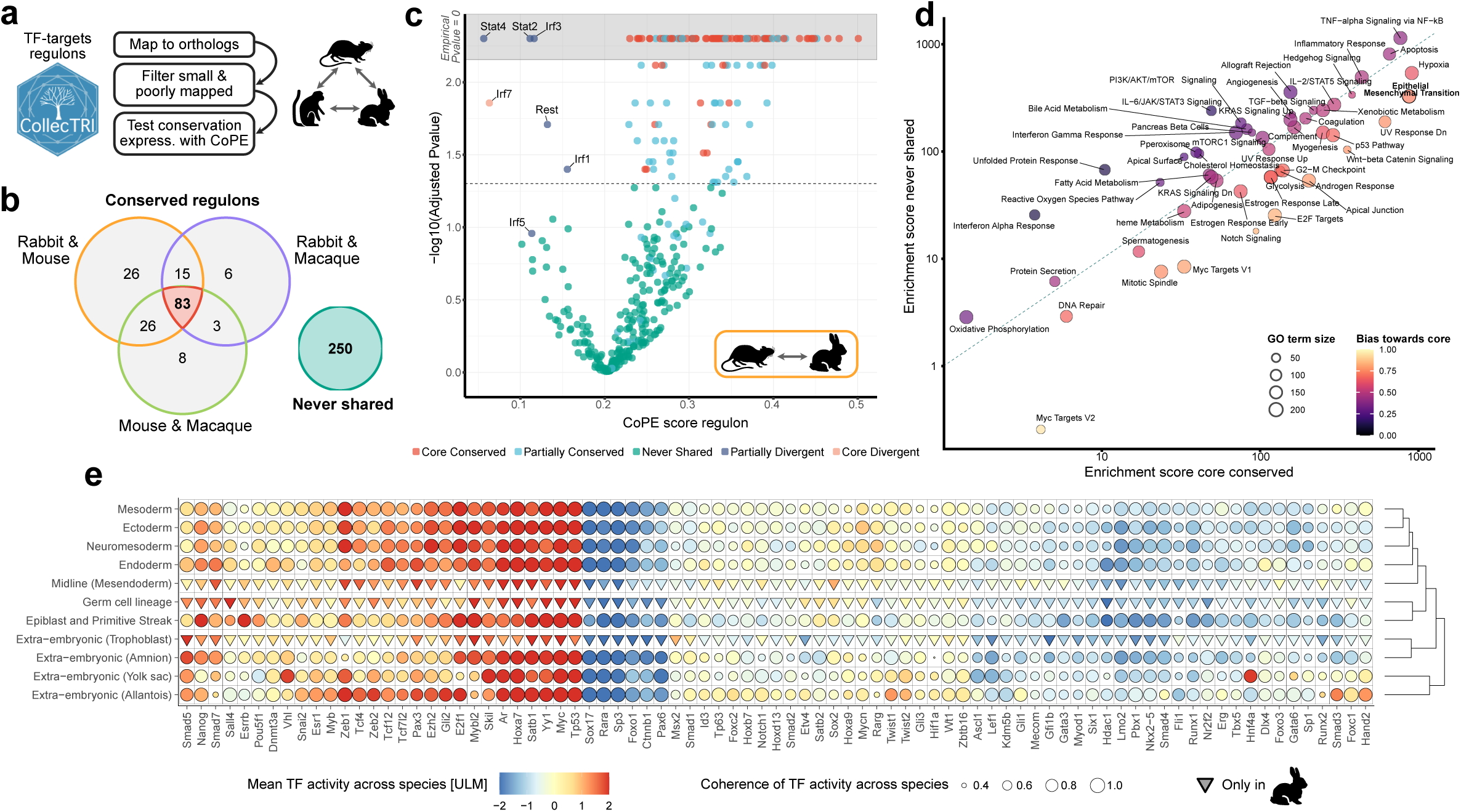
Identification of conserved transcription factor programs across mouse, rabbit, and crab-eating macaque. **a**, Quantification of regulon conservation. Regulons from Collectri (i.e. TFs and their target genes) were mapped to orthologs common to the three species. Regulons that were small or poorly mapped were filtered out. Regulons were given a CoPE score by averaging the individual CoPE scores of their target genes. **b,** Number of Collectri regulons found to be conserved across species pairs. Core TFs are highlighted in red. **c**, Details of the selection of statistically significant genes for the mouse-rabbit pair based on regulon CoPE scores. P-values were derived through random sampling of gene groups of the same size as individual regulons. The grey box indicates empirical p-values equal to 0. The regulon categories were determined by considering all pairs of species. Core conserved (resp. divergent) regulons were found to be significantly more (resp. less) conserved than expected across all pairs of species. **d**, GSEA with the human MSigDB Hallmarks terms^35,36^. Enrichment scores for each term were calculated by pooling all the regulons from the core conserved set, and all the regulons from the never shared set independently. Enrichment scores were calculated using EnrichR’s^37^ combined enrichment scores. **e**, Core conserved regulon activity, averaged by harmonized tissue annotations. Regulon’s activity was evaluated with Univariate Linear Model (ULM)^38,39^ in every neighbourhood independently and averaged by harmonized tissue annotation across species. The harmonized annotations were obtained by manually grouping matching cell types; tissues observed in rabbit only are represented with inverted triangles. The coherence of TF activity was derived by independently ranking the regulons’ activity in each species, and taking the difference between the maximum and the minimum rank across the three species.

As expected for one-to-one orthologs, we found no regulons with a significant negative CoPE score, and only 7 with a low CoPE score (∼[0, 0.10]) indicating greater divergence in expression between species (Fig. 4c, Supplementary Table S2). Among them, only one regulon (Irf3) was shared across all pairs of species, but most were closely related to interferon signalling (Irf1, Irf5, Irf7, Stat2, Stat4). The immune system is known to be both highly diverged and also unlikely to play a critical role in cell type specification during gastrulation as it is still forming. Interestingly, the RE1-silencing transcription factor (Rest) regulon, a transcription factor involved in the repression of neuronal gene expression in non-neuronal tissues, was also found to be less conserved in the rabbit-mouse and rabbit-macaque pairs, but not in the mouse-macaque pair, indicating that the role of this regulon (or the timing at which it is active) might differ in the rabbit. Surprisingly, we found that the majority of regulons were not significantly more conserved than random samples of orthologs. This result suggests that evolutionary pressure on expression levels may be strong only for a select few developmental processes. Finally, we identified a set of 83 TFs, which we refer to as core TFs, whose regulons were significantly more conserved across all species pairs, suggesting a critical role in gastrulation.

We characterized the function of conserved and less conserved regulons by performing GSEA on the genes that compose them (Fig. 4d). This revealed that the less conserved regulons were enriched in terms related to immunity, such as interferon alpha or Jak/Stat pathway. In contrast, the core regulons were enriched in terms related to the cell cycle (Myc Target, E2F targets, glycolysis) and epithelial-to-mesenchymal transition (EMT, Notch, Wnt, apical junction). This finding highlights a very strong conservation of the mechanism defining the three germ layers, and suggests that a multitude of TFs are required for it. The small overlap between the targets of the core TFs (Fig. S5) also implies that this set of TFs must be working together with less-conserved TFs to achieve a quantitative regulation of the target transcripts. Among the core TFs, we also recovered several highly conserved families, including all Hox genes responsible for antero-posterior patterning that met our gene set size criteria. This further highlights the conservation of processes defining the body plan.

Finally, we characterized the tissues in which the core TFs were active, and assessed the coherence of their activity across cell types and species (Fig. 4e). For this, we computed the activity of the core TFs in individual neighbourhoods using the Univariate Linear Model (ULM) method^38,39^. This method estimates activity by quantifying the degree to which the expression of a TF’s target genes exceeds that of non-target genes. We then aggregated their activity at the level of manually harmonised germ layer annotations, and averaged TF activity within each germ layer across the three species. We found that the core TFs displayed a large range of activity patterns: from very active TFs to very lowly active (yet still conserved) across all germ layers, and a few specialised TFs displayed activity in specific tissues. Additionally, we observed that the spread of TFs activity was very consistent across species, as expected for conserved processes.

### Augmentation of sparse atlases, cross-species prediction, and calibration across biological models with RIMA matching

Despite a significant drop in sequencing costs, generating single-cell atlases remains costly, and access to embryonic material is limited by both ethical considerations and availability. These limitations can lead to the creation of sparse atlases – with missing or poorly represented tissues or time-points – and increase reliance on *in vitro* models, for which the molecular correspondence to *in vivo* counterparts is not always well established.

To address these challenges, we tested RIMA’s capabilities to impute missing profiles in sparse atlases and evaluated whether its neighbourhood mapping could be used to capture and adjust for expected expression differences between a model system and its biological target. For this test, we withheld the pairs of neighbourhoods that include data from the 8th embryonic day of the mouse (i.e. sampled from embryos aged E8.0, E8.25, E8.5 and E8.75; collectively referred to as E8), a day marked by the rapid transition from undifferentiated germ layers to differentiated cell lineages (Fig. S4). Using the pairs from the remaining time-points, we trained a regressor to predict mouse expression profiles from their matched rabbit counterparts. Finally, we imputed the withheld E8 mouse neighbourhoods by applying the regressor to their matched rabbit neighbourhood and evaluated the prediction accuracy (Fig. 5a,b, Fig. S6b, Supplementary Table S3).

**Figure 5:**
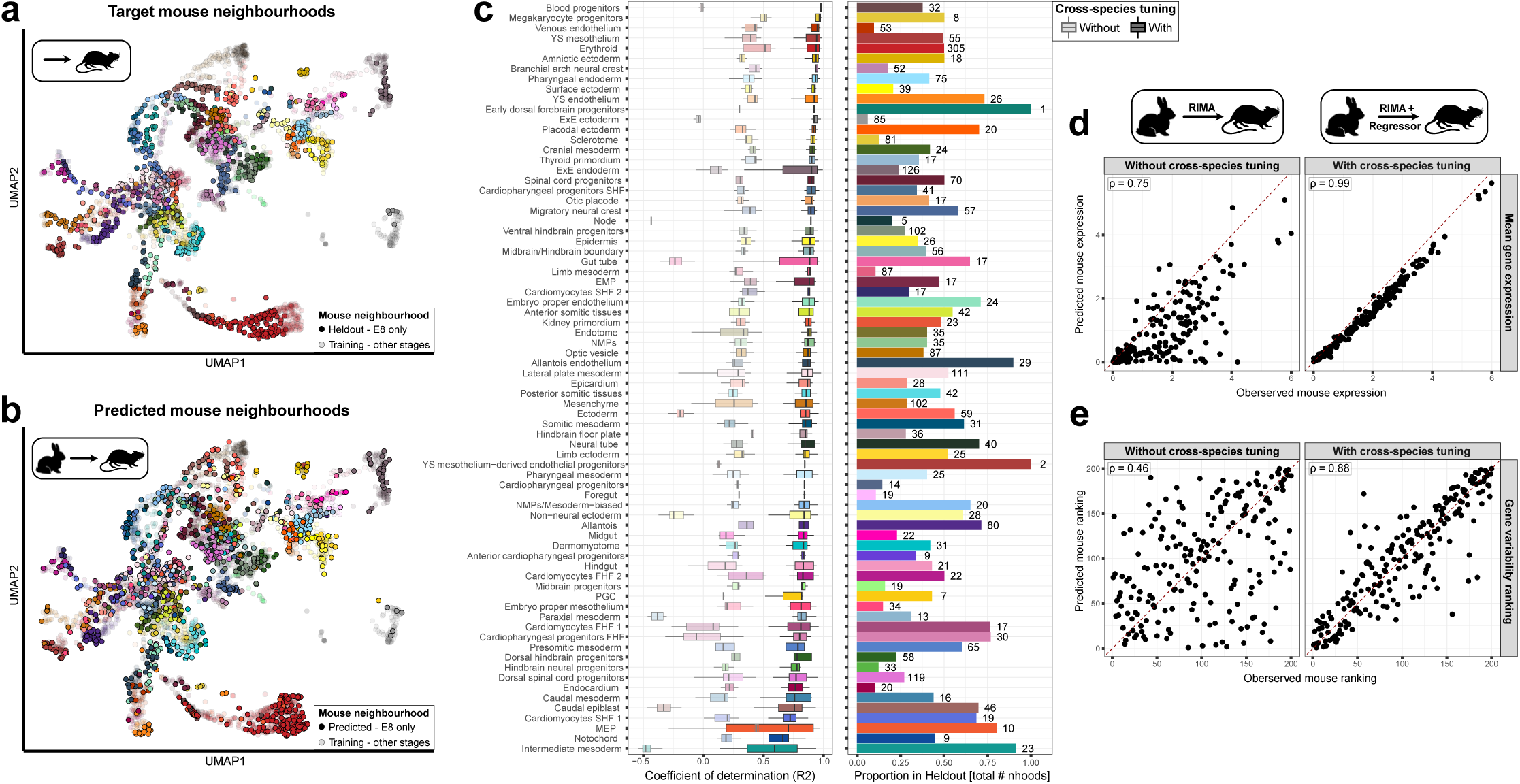
Cross-species prediction of missing mouse transcriptomic profiles. **a-b**, UMAP projection of the mouse atlas, batch-corrected with fastMNN, integrated together with neighbourhoods predicted from rabbit profiles. The prediction training set was formed with the mouse-rabbit neighbourhood pairs where the mouse neighbourhood was coming from E6, E7, and E9 embryos. The observed E8 target mouse neighbourhoods (top) and the E8 mouse neighbourhoods predicted from rabbit (bottom) are represented in solid colours. **c**, Left: quality of fit of held out neighbourhoods, approximated using their matched rabbit neighbourhoods without (transparent) or with (solid) transformation with a random forest. Right: proportion of neighbourhoods coming from E8 mouse embryos forming the holdout set, the number of withheld neighbourhoods are indicated next to the bars. **d-e**, Observed gene mean logcounts (top) and gene variability rank (bottom) in E8 mouse neighbourhoods against neighbourhoods approximated using matched rabbit neighbourhoods before (left) or after (right) transformation with a trained random forest. Pearson correlations are indicated.

In practice, we trained a random forest regressor tasked with minimizing the L2 difference between the normalized logcounts of mouse neighbourhoods and the values predicted from the matched rabbit logcounts (Fig. S6b). To facilitate the training, while retaining biological information, we limited the task to a panel of 200 genes returned by GeneBasis, which captured most of the mouse atlas connectivity (Fig. S6c). We also implemented a data augmentation strategy, randomly dropping cells from training rabbit neighbourhoods to increase the number of training examples and improve prediction stability (Fig. S6a). Finally, we integrated the predicted mouse neighbourhoods into the mouse atlas with fast Mutual Nearest Neighbour (MNN)^42^, treating them as a single batch (see Methods, Fig. 5b).

We found that the random forest successfully generated mouse-like neighbourhoods that could be embedded within the original mouse atlas (Fig. 5a,b). With the exception of megakaryocyte-erythrocyte progenitors (MEPs), the predicted profiles were systematically better at recapitulating the missing mouse profiles than the source rabbit neighbourhoods that were used as starting points for the predictions (Fig. 5c). This gain of quality was present even for cell types where a large fraction of the neighbourhoods were withheld (Fig. 5c) (e.g. intermediate mesoderm), and remained consistent in more divergent tissues such as the extra-embryonic tissues (e.g. yolk sac). Importantly, the closer fit of predicted tissues after cross-species tuning was not only due to correcting the average expression of each gene (Fig. 5d), but also came from a restoration of the variability in expression across tissues. This can be seen from the ranking of genes by variability, which was much closer to the mouse ranking in the predicted mouse neighbourhoods than in the source rabbit neighbourhoods (Fig. 5e).

Altogether, these results indicate that RIMA can leverage partial atlas matching to generate realistic and accurate atlas augmentations, as well as correct deviations between a model and its target for out-of-sample predictions, opening the door to better translation from *in vitro* to *in vivo*, and to a robust augmentation of sparse atlases.

## Discussion

In this study, we present a comparative analysis of gastrulation and early organogenesis between three mammals and highlight both high-level and subtle (dis)similarities between their transcriptomic profiles. Central to our analysis, we introduced RIMA, a versatile matching method for single-cell transcriptomics that enables a range of downstream comparative tasks at the scale of cell neighbourhoods. Working at this scale not only provides technical advantages such as speed and scalability, but also allows for a quantitative, fine-grained comparison of the input atlases by locally matching cell identity. Operating at near-single-cell resolution with matching states means that the comparison controls for confounding factors, such as batch effect and cell type composition. Such comparison of biological states on a “like-for-like basis”, highlights subtle similarities and differences while avoiding the signal loss induced by techniques such as pseudo-bulking or over-reliance on manual annotations.

Our cross-species analysis shows that early development of mammals, albeit heavily conserved overall, displays varying degrees of molecular similarity across tissues and species. As previously reported, we found that extra-embryonic tissues can be reliably identified across species but are, on average, transcriptionally more distinct than tissues from the embryo-proper^6^. We also recapitulated the traditional model of the developmental hourglass and located a similarity bottleneck around E7.75 and GD8 in the mouse and the rabbit, respectively. Importantly, our analysis identifies the hourglass pattern based on molecular similarities independently of anatomical features or cell type annotations. We found that the hourglass timing is likely explained by the conservation of the processes giving rise to the future body plan, notably the migration and differentiation of early tissues into the germ layers through epithelial-to-mesenchymal transition. The presence of such a molecular similarity bottleneck, despite differences in embryo shapes and speed of development, highlights that signalling pathways have evolved to robustly carry consistent signals in presence of large species-specific traits such as the distinct cup-like morphology of the mouse embryo^6^.

Beyond broad descriptive comparison, RIMA’s matching enabled us to uncover underlying molecular mechanisms driving similarity through the identification of conserved processes at multiple scales. At the whole-embryo level, we identified a core set of TF programs (regulons) common to the mouse, rabbit, and macaque. Many of these core TFs are related to the EMT, a key process for the formation of the mesoderm and endoderm around E7.5-E7.75 in the mouse. This timing coincides with our estimated window for the hourglass bottleneck, and highlights the strong evolutionary pressure on the formation of the body plan across species. In another study^4^, a similar mouse-rabbit analysis identified TFs with reciprocally correlated expression across matched cell states. Surprisingly, the two approaches yielded largely distinct conserved TF lists: only 20 of 156 TFs identified in our study overlapped with the 76 from the other. Part of this difference reflects the limitations of our approach which relies on prior knowledge: 5 of the non-recovered TFs were absent from Collectri, and 24 lacked sufficient one-to-one orthologs to meet our regulon size threshold. The discrepancy between our approach that relies on target genes’ expression rather than on TF expression reflects the complexity of transcriptional regulation during development and calls for a deeper investigation, notably by extending to other modalities (e.g. proteomics) to understand how the multiple sources of regulation contribute to the course of development in each species.

RIMA also enabled a more focused comparison of erythrocyte formation, where we identified conserved transcriptional boosts of key developmental genes involved in erythrocyte development^34^. However, we found that the sequence and the onsets of boosting were subtly different between the mouse and the rabbit. How much flexibility is possible around these dynamics to reach functional haematopoiesis – such as the precise sequence, timing, or amplitude of boosting – remains unclear. Further experiments could help clarify these aspects and verify if, for example, the sequences of boosting are interchangeable between species or have emerged as species-specific requirements during evolution. This analysis provides insight into potential flexibility of haematopoiesis emergence, which could help to derive better *in vitro* models and produce blood cells more faithful to their *in vivo* counterparts, with potential implications for translational research.

Although workflows that rely on integration or foundation models are becoming more interpretable^43^, we demonstrated that comparative approaches, such as RIMA, provide a simpler, flexible alternative for a wide range of downstream tasks with minimal assumptions about the data. While we applied RIMA here to cross-species transcriptomics, its framework is readily applicable to same species comparisons, expanding its potential applications (e.g. *in vitro* vs *in vivo*, healthy vs disease groups). Building upon our three-way comparison between mouse, rabbit, and macaque, we also envision meta-analysis as an important avenue of development for RIMA, enabling the pairing of matching cell states across studies and supporting controlled comparisons with improved statistical power.

Finally, we demonstrated that RIMA’s application can go beyond characterizing differences between systems and can also be used to address limitations of existing experimental designs. For example, while model organisms are often used as “drop-in approximations” for their target, we found that even after carefully matching rabbit to mouse cell states, a large gap remains between their transcriptomics profiles. However, we show that a proper matching of cell states provides a solid basis to refine this use of biological models and generate cross-system predictions that restore the target biological signal. In our case study, we could map observations from the rabbit to the mouse and generate accurate imputation of gastrulation profiles that accurately reflect mouse biology. We envision that such predictions can provide *in silico* estimation of the cell states of expensive or hard to obtain biological material (e.g. human developmental data), enabling sparse atlases augmentation, and the derivation of better *in vitro* models, with implications for basic and translational research.

## Methods

### Generation and processing of scRNA-seq atlases

The processed versions of the mouse (*Mus musculus*)^22^ and the rabbit (*Oryctolagus cuniculus*)^6^ scRNA atlases were used throughout this study. These “ready-to-be-used” versions were made available in the original publications. The mouse and rabbit atlases are respectively available at: https://marionilab.github.io/RabbitGastrulation2022 and https://marionilab.github.io/ExtendedMouseAtlas. The authors followed a consistent preprocessing pipeline to generate both atlases. Both atlases were produced using the 10X Genomics Chromium system with chemistry kit version 3, except for a subset of the mouse atlas which was generated using version 1. Further details are available in the original publications.

We have regenerated the crab-eating macaque (*Macaca fascicularis*) atlas^7^ from the raw read files to make it consistent with the other two atlases and to use a more recent version of the genome annotation than the one that was used in the original publication. We first downloaded the fastq files related to the 7 samples and aligned them with CellRanger (v7.1) to a reference atlas which we built with Ensembl’s genome and annotations (Macaca_fascicularis_6.0, Ensembl release 108). We then preprocessed the counts consistently with the pipeline used for the mouse and rabbit atlases. We performed initial quality controls to remove “noisy cells” using the adaptive thresholds of the *scater*^44^ R package (library size > 1432, number of features > 1330, mitochondrial content < 3.83%). The reads were normalized with the “deconvolution method” using the *calculateSumFactors* function from the *scran*^45^ R package, and doublets were removed from each sample independently using the *cxds_bcds_hybrid* function from the *scds* R package^46^. We then integrated the samples using the *fastMNN* function from the *batchelor* R package^42^ with parameters: 3000 highly variable genes, 50 principal components, and 20 neighbours. We integrated the samples from oldest to youngest embryos and integrated the largest samples first for embryos of the same age. Upon inspection of the integrated cells, we identified and discarded a poorly integrated cluster of cells, all coming from the same sample (Carnegie Stage 8 - embryo 1) which was originally discarded by the authors and likely composed of doublets missed in the previous step with our looser QC parameters. We ran a new round of integration with fastMNN after dropping these cells, using the same parameters as earlier. We kept the cell annotations from the original publication where possible, and gave a cell type annotation to the cells unique to our pipeline with a KNN classifier using their 9 closest neighbours in the annotated cells. The number of neighbours was tuned by maximizing the F1 score of KNN classifiers trained and tested on the already annotated cells. The final atlas was composed of 49,970 cells and 21,073 genes. A detailed description along with the code to regenerate this atlas is available here: https://github.com/ma-jacques/macaque_atlas.

### Neighbourhoods definition

We defined neighbourhoods independently for each atlas using a refined sampling strategy^23,24^. This strategy starts by randomly picking a set of cells, and defines a neighbourhood around each of these index cells by taking their neighbouring cells in a KNN graph. The neighbourhoods formed this way are then recentered around their average cells, which become the new neighbourhood index cells, and their KNN neighbours form the final neighbourhood.

To define neighbourhoods in our 3 atlases, we considered independent KNN graphs that were built on the integrated PCA coordinates produced by fastMNN for each atlas. We used the Euclidean distance in this set of coordinates to identify the nearest neighbouring cells around index cells. We used the functions *buildGraph* and *makeNhoods* from the *miloR* R package with the following parameters for the mouse, rabbit, and macaque respectively: 30 nearest neighbours to define the KNN graphs; 75, 50, 50 MNN-corrected PCs to calculate the Euclidean distance between cells; 1%, 5%, 5% of the total number of cells to randomly sample the initial index cells. This formed 3448, 5254, and 2063 neighbourhoods for the mouse, rabbit, and macaque respectively.

The neighbourhoods were annotated (e.g. cell type, developmental day) using the most frequent label of the cells which composed them.

### Cross-species neighbourhoods matching with RIMA

RIMA’s matching relies on the calculation of similarity of cell neighbourhoods between species. The similarity between two neighbourhoods is defined as the Spearman correlation between their average gene expression levels. The calculation of all pairwise similarities between neighbourhoods defines a weighted bipartite graph, connecting the neighbourhoods between input species with undirected edges that carry the Spearman correlation of the neighbourhood-neighbourhood pairs.

#### Identification of the gene panel used to calculate the neighbourhood-neighbourhood similarity

To enable the calculation of the Spearman correlation across neighbourhoods of different species, we first established a correspondence of the genes across the mouse, rabbit, and macaque, by mapping the genes to their one-to-one mouse orthologs, when available. We then used the intersection of the mapped and observed genes across species pairs to compute the neighbourhoods’ similarity. We identified two different sets of genes to use for cross-species matching with RIMA: a first set with the mouse and rabbit only (Fig. 2, 3, 5) and a second set where we added the macaque atlas (Fig. 4). We used the common one-to-one orthologs observed in the mouse and rabbit atlases for the first set, and the common orthologs to the mouse, rabbit, and macaque for the second set. We identified one-to-one orthologs with Ensembl’s annotations, which were accessed through the *biomaRt* R package^47^. This resulted in a set of 13,968 orthologs to compare the mouse and the rabbit, and a set of 12,121 orthologs to compare the mouse, the rabbit and the macaque.

#### Significance of the similarity of a neighbourhood-neighbourhood pair

We wrote an R implementation of RIMA to match neighbourhoods across atlases available at https://github.com/ma-jacques/RIMA. We defined the similarity between pairs of neighbourhoods across species with the Spearman correlation of the average normalised log counts of the orthologs, obtained as described earlier. We calculated the neighbourhoods’ average expression with the *calcNhoodExpression* function from the *miloR* R package, and the Spearman correlation with the *cor* R function.

We then used RIMA’s resampling strategy to define the significance of the directed edges connecting two neighbourhoods. For this, we compared the correlation between original neighbourhoods to correlations between original and resampled neighbourhoods, for which cell identities have been randomly resampled. The random resampling of the neighbourhood’s cells was done by keeping one atlas intact and ressampling the other one, and repeated in the other direction. This yielded two measures of significance for each edge connecting two neighbourhoods, calculated as:

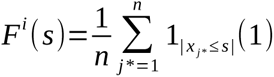

Where (1) is calculated for each original neighbourhood (*i*) from the non-resampled atlas, and represents the empirical cumulative distribution function (*F*) of the Spearman correlation (*s*) between the original neighbourhood *i* and the *n* resampled neighbourhoods (*j\**) from the other atlas. This is then used in:

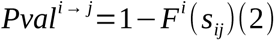

Where (2) represents the p-value associated with the directed edge that links the original neighbourhoods *i* to *j* and carries the Spearman correlation *s_ij_*. For each edge, the p-value in the opposite direction *Pval ^j→i^* is also calculated by resampling the other atlas and estimating *F ^j^*(*s*). The combined p-value of the undirected *i↔ j* edge is obtained by combining both directional p-values *Pval^i→^ ^j^*and *Pval ^j→i^* with Simes’ method, a combination method that is robust to positively dependent values^26,48^. In our comparisons, the combined p-values were adjusted with Holm’s method for family-wise error rate, and edges with a p-value below 0.05 were deemed significant, and considered as candidate matches for the next step.

In this study, we resampled the cell identities of each atlas 10 times, keeping the same structure of neighbourhood graphs (i.e. the number and size of neighbourhoods were kept intact). To produce truly unspecific neighbourhoods, rather than permutation, we used a weighted resampling scheme with replacement, where each cell was given a probability of sampling that was inversely proportional to the frequency of its cell type. This ensured that the permuted neighbourhoods did not closely resemble the original neighbourhoods due only to the very large imbalance of cell type composition in the atlases (e.g.,, in the mouse atlas 38,975 cells were annotated as erythrocytes, while only 190 cells were annotated as early dorsal forebrain progenitors). Without weighted resampling, this imbalance would bias the resampled neighbourhoods to resemble erythrocytes, hence the matching would be more stringent for cell types that resemble them.

### Matching significant neighbourhood-neighbourhood pairs

In the previous step, RIMA trims the insignificant neighbourhood-neighbourhood edges in the weighted bipartite graph formed by calculating the neighbourhood’s pairwise across species. This trimmed graph is finally resolved into a one-to-one matching of neighbourhoods by finding a matching that maximizes the sum of the weight of the edges in the matching. For this, we used the implementation provided by the function *max_bipartite_match* in the *igraph* R package^49^.

### Correlation of paired expression (CoPE) score

We define the correlation of paired expression (CoPE) score as a measure of conservation of gene expression levels across species. Let *G*=(*U, V, E*) be the graph built by RIMA to match the atlas *U* to the atlas *V* with edges *E*. The edges represent the *n* neighbourhood-neighbourhood matches (*u_i_, v _j_*) which connects *u_i_ ɛ U* and *v _j_ ɛ V* between the two atlases. The nodes *u_i_* and *v _j_* carry a range of numerical features, representing the average gene expressions of the cells in these neighbourhoods. Let *f _i, k_* and *f _j, k_*denote the average normalised log count of gene *k* in the neighbourhoods *u_i_* and *v _j_*, respectively.

The CoPE score of the gene *g* is defined as: *C_g_*=*S* (*f _i, g_, f _j, g_*); where *S* is the Spearman correlation function.

We extend the CoPE score of individual genes to Collectri^39^ regulons by calculating the CoPE score for each gene in the regulon (including the controlling TF) and averaging the CoPE score of all genes in the regulon.

### Comparison of erythrocyte development between the mouse and rabbit

#### Identification of cells from the erythrocyte lineage

The cells belonging to the erythrocyte lineage were independently identified in the mouse and the rabbit atlases, from early undifferentiated tissues to differentiated erythrocyte cells. We used a freshly downloaded version of the processed atlas, as provided by the original studies. We used all available genes in these atlases, including cycling genes and genes without orthologs. For the mouse atlas, we excluded cells coming from batches where embryos from multiple stages were mixed together.

We first made a loose selection of the lineage by manually selecting the cells annotated with a cell type known to belong to the lineage, or sharing common progenitors:

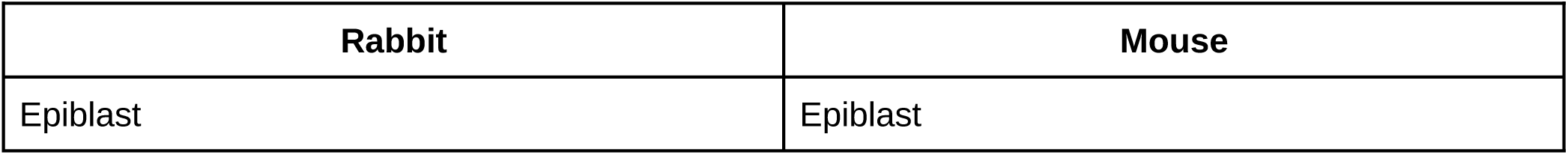
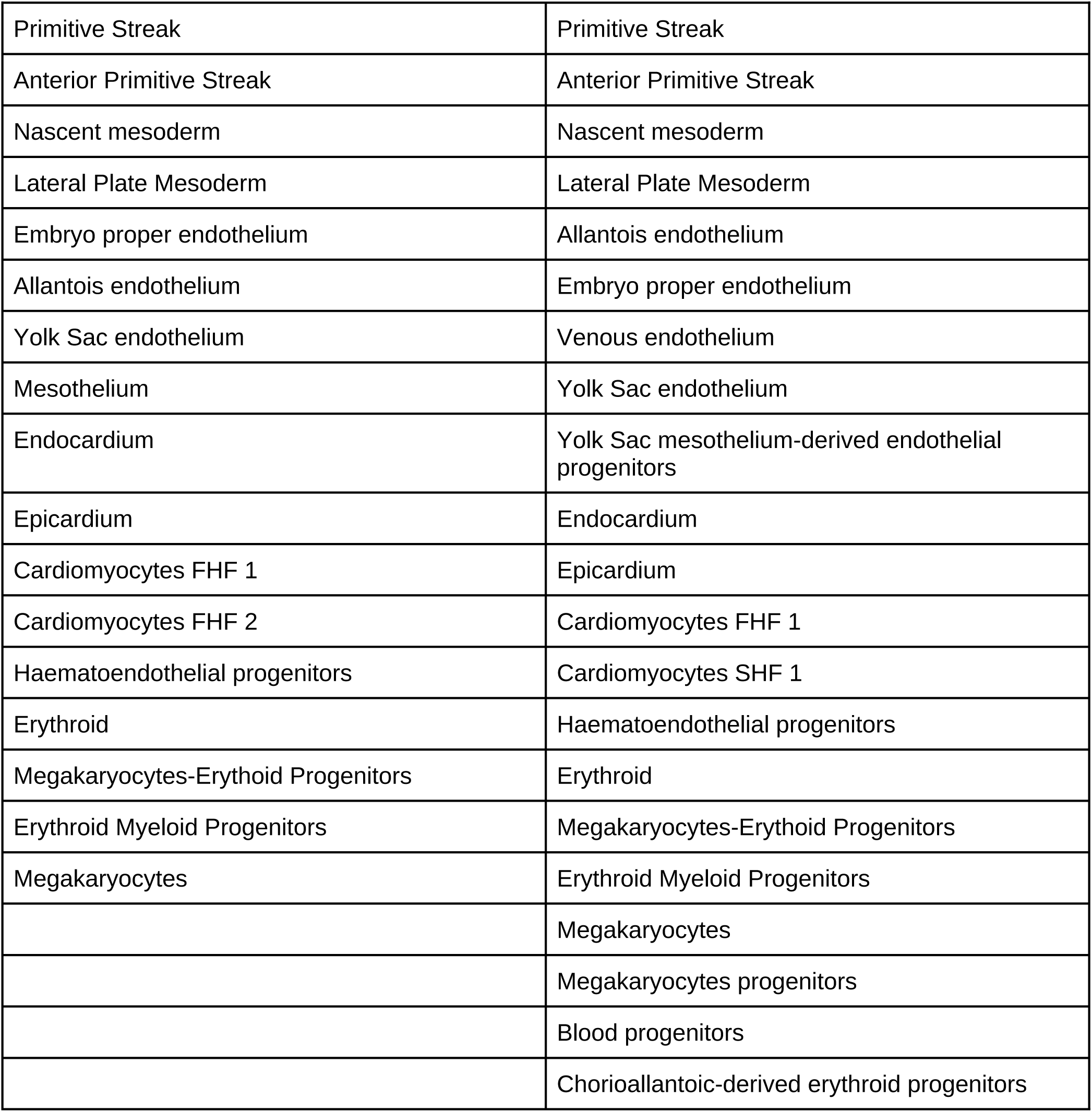

| Rabbit | Mouse |
| --- | --- |
| Epiblast | Epiblast |
| Primitive Streak | Primitive Streak |
| Anterior Primitive Streak | Anterior Primitive Streak |
| Nascent mesoderm | Nascent mesoderm |
| Lateral Plate Mesoderm | Lateral Plate Mesoderm |
| Embryo proper endothelium | Allantois endothelium |
| Allantois endothelium | Embryo proper endothelium |
| Yolk Sac endothelium | Venous endothelium |
| Mesothelium | Yolk Sac endothelium |
| Endocardium | Yolk Sac mesothelium-derived endothelial progenitors |
| Epicardium | Endocardium |
| Cardiomyocytes FHF 1 | Epicardium |
| Cardiomyocytes FHF 2 | Cardiomyocytes FHF 1 |
| Haematoendothelial progenitors | Cardiomyocytes SHF 1 |
| Erythroid | Haematoendothelial progenitors |
| Megakaryocytes-Erythroid Progenitors | Erythroid |
| Erythroid Myeloid Progenitors | Megakaryocytes-Erythroid Progenitors |
| Megakaryocytes | Erythroid Myeloid Progenitors |
|  | Megakaryocytes |
|  | Megakaryocytes progenitors |
|  | Blood progenitors |
|  | Chorioallantoic-derived erythroid progenitors |

We then used *CellRank 2*^50^ (v2.0.6) to estimate the fate of these cells. We calculated two cell-cell transition matrices using the RealTime kernel and the CytoTrace kernel, and combined them with an equally weighted sum. For both kernels, we used a KNN graph built with *Scanpy’s*^51^ *pp.neighbor* function, using the MNN-corrected PCA space provided by the original authors.

Unless specified hereafter, we used the default parameters for the estimation of cell-cell transition matrices. For the CytoTrace kernel, we preprocessed the data with *scVelo’s*^52^ *pp.moments* function using 30 PCs and 30 neighbours, and calculated the transition matrix with a soft thresholding scheme with parameter nu = 0.5. For the RealTime kernel, we preprocessed the data with *Moscot’s*^53^ TemporalProblem (v0.4.0) using the mouse proliferation and apoptosis gene sets, and temporally linked the cells using the cell’s embryonic day as timestamp; we then computed the transition matrix with the ‘all’ transition logic, and weighted the connectivities’ self transition with a weight of 0.2.

Using the combined transition matrix and the Generalized Perron Cluster Cluster Analysis (GPCCA) estimator, we identified 10 and 20 macrostates in the mouse and the rabbit, respectively. Among the macrostates, we manually chose a single state to best represent the following terminal nodes: megakaryocytes, yolk sac endothelium, erythroid and cardiomyocytes. We also selected the most representative epiblast and primitive streak macrostates as initial states.

In both species, we estimated the fate probability for each terminal state with the GPCCA estimators, and selected the cells that were likely to have an erythroid fate. To do so, we selected the cells whose erythroid fate probability was above 42%, and 9% for the mouse and the rabbit, respectively. These thresholds were determined by manual inspection of the selected cells, notably by ensuring that all cardiomyocytes were excluded from the selection. For both species, we annotated a neighbourhood as belonging to the erythroid lineage if it was composed of at least 75% of cells from the erythroid lineage.

#### Pseudotime ordering of the erythrocyte lineage

After selecting the cells that were likely to have an erythrocyte fate, we ordered them along pseudotimes that represented the progression from undifferentiated tissues (epiblast, primitive streak), to differentiated erythrocytes with *Palantir*^54^ (v1.3.6).

To build the pseudotime axis, we first found its root by identifying a start cell in each species using the CytoTrace score as a proxy to measure the degree of differentiation. The start cells were defined from the cells annotated as epiblast by choosing those with the smallest CytoTrace score in each species.

We then used Palantir to build the pseudotime for each species on 10 diffusion components derived from the MNN-corrected PCs of the selected cells, and used the start cell to define the orientation of the pseudotime axis. Finally, we normalized the pseudotime in both species between 0 and 1, where 0 represents the most undifferentiated epiblast cells, and 1 represents the most differentiated erythrocytes.

#### Construction of MURK gene expression time-series and logistic function fitting

We constructed a time-series of expression of the MURK genes in the rabbit and the mouse representing the expression of these genes as cells progress along the erythrocyte differentiation trajectory. For this, we took the average log counts of these genes in the species’ neighbourhoods, and placed them on the erythrocyte trajectory by taking the average of the normalised pseudotime of the cells which composed the neighbourhoods. To express the time-series on a common pseudotime coordinate, we used RIMA’s neighbourhood pairs to convert the pseudotime of rabbit neighbourhoods into the pseudotime of their matched mouse neighbourhood.

To characterize the transcriptional boost of the MURK genes during erythrocyte differentiation, we considered only the neighbourhoods whose mouse pseudotimes were in the range [0.65, 1]. This range corresponds to the transition from haematoendothelial progenitors to erythroid-myeloid progenitors, where most MURK genes start to be expressed. We then normalized each gene’s expression between 0 and 1 on this segment, and smoothed the resulting time-series with a rolling mean of length 20 to facilitate the fitting of a logistic (sigmoid) function. The sigmoid was fitted using the *nls* function in R with the formula:

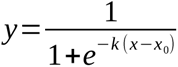

Where *y* represents the expression of a MURK gene, *x* represents the mouse pseudotime, *k* represents the steepness of the curve, and *x*_0_ represents the midpoint of the sigmoid. The fit of each MURK gene time-series was allowed a maximum of 50 iterations, with starting values *x*_0_=0.5 and *k* =1.

### Identification and analysis of conserved processes of the erythrocyte lineage across mouse and rabbit

We have characterized well-conserved genes along the development of erythrocytes through gene set enrichment analysis (GSEA) (Fig. 3). For this, we first selected the genes to analyse by keeping genes which displayed both a high CoPE score, and a high level of expression. The CoPE scores were calculated by computing the Spearman correlation of gene expression between the mouse and the rabbit, only considering the pairs of neighbourhoods (as returned by RIMA) where both the mouse and the rabbit neighbourhood were composed of at least 75% of cells coming from the erythrocyte lineage (see earlier).

The CoPE score cutoff was determined with a bootstrap procedure where genes’ identities were randomly permuted in both species independently. The CoPE score cutoff was set to keep only the genes whose score was greater than 99% of the scores obtained by bootstraping the data. The high expression cutoff was set to only keep the genes that displayed a raw count greater than 2 in at least one neighbourhood among all the erythrocyte neighbourhoods.

We performed a GSEA on the selected genes using the *enrichR*^37^ R package, using the 2023 GO biological processes gene sets^32,33^. The terms whose adjusted p-value was below 0.05 were considered significant, and visualised with the *GO-figure!* package^55^, using a go-term similarity threshold of 0.6.

### Identification of core Collectri regulons across mouse, rabbit, and macaque

We identified TF regulons which are conserved across mouse, rabbit, and macaque among the mouse Collectri^39^ gene regulatory network. As a first step, we filtered the regulons to keep only those where at least 10 genes could be mapped to a one-to-one mouse ortholog. Additionally, we filtered out regulons where less than 50% of the original regulon genes could be mapped. Out of the 1072 original regulons, 424 passed these filters and were analysed further.

We expanded the notion of CoPE score from individual genes, to the scale of regulons, by taking the average CoPE score of the TF and all target genes that form a regulon. We then assessed the statistical significance of each regulon’s CoPE score by randomly sampling 1000 pools of genes (among the one-to-one orthologs) of the same size as each regulon, and averaging the CoPE scores of the genes in these random samples. Empirical two-sided p-values were derived by comparing the regulon’s CoPE score to these random samples with the following procedure:

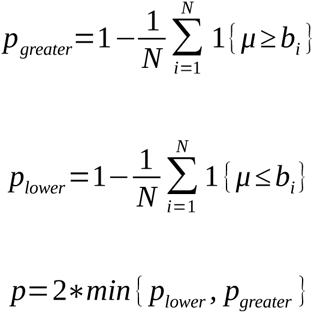

Where *μ* is the regulon’s CoPE score, compared to N random gene samples of same size as the regulon which carry the CoPE scores {*b*_1_ *,b*_2_, … *, b_N_* }. The regulon was called conserved if *p_lower_ ≥ p_greater_*. Finally, the regulons with an FDR-adjusted p-value below 0.05 were called significantly conserved/less conserved, depending on the effect direction. CoPE scores were calculated and assessed for all species’ pairs, and we defined core regulons as those that were significantly conserved across all species’ pairs.

We performed a GSEA of regulons (i.e. the TF and all its target genes) using the *enrichR*^37^ R package, using the 2020 human MSigDB Hallmark gene sets^35,36^. We performed 2 separate rounds of GSEA, one with the core conserved regulons, and one with the regulons that were not shared between any pair of species. For each round we extracted the “combined score” of each term as returned by *enrichR*, and used them to define the term’s enrichment score in both sets of regulons. A “bias towards core” for each term was defined as the ratio between the enrichment score in core regulons, over the sum of the enrichment scores in core and never shared regulons.

The core regulons were further characterized by calculating their TFs’ activity in individual neighbourhoods using the Univariate Linear Model (ULM) method from the *decoupleR* R package^38,39^. To obtain a more faithful estimation of the regulon’s activity in each species, we used all available genes of the regulons in each species, not only the genes that could be mapped to a one-to-one mouse ortholog. TF activity was then averaged at the level of tissues by averaging the ULM score of the neighbourhoods, following their cell type annotations. Tissue annotations were manually defined to harmonise cell type annotations across species.

### Cross-species prediction

For brevity, we define hereafter the expression profile of a neighbourhood, as the average of the normalised log counts of the cells that compose the neighbourhood. Using RIMA’s neighbourhood-neighbourhood pairs across mouse and rabbit, we established the input-output training pairs for a random forest regressor. This model was tasked with predicting mouse neighbourhood expression profiles from their paired rabbit neighbourhood expression profiles by minimising the L2 distance between them. The random forest was trained with the *scikit-learn*^56^ (v1.4.1) Python (v3.11) package, using the *RandomForestRegressor* function with default parameters. Most notably, we used 100 trees and did not limit the depth of the trees.

To simulate missing data in the mouse atlas, we withheld all neighbourhoods annotated with any embryonic day 8 time point (E8.0, E8.25, E8.5, E8.75) (Fig. S6b), using the remaining neighbourhoods as the training dataset. To prevent information leakage between the training and held-out sets — which could inflate predicted performance and bias evaluation — we implemented several steps. First, we completely excluded the neighbourhoods annotated as “Mixed gastrula” both from the training and the held-out set, as we could not trace back the age of the cells that composed them. Then, as the cells composing neighbourhoods typically do not all come from the same time-point, we also removed any E8 cell from the training neighbourhoods prior to calculating their expression profiles. Conversely, we also removed all non-E8 cells from the heldout neighbourhoods. We then calculated a set of metrics to ensure that removing the cells did not affect the neighbourhood’s average logcounts (Fig. S6d).

We also implemented a data augmentation strategy (Fig. S6a) to increase the number of training examples and encourage the learning of a function that is robust to small variations. This was also motivated by the observation that the number of neighbourhoods coming from the different cell types was extremely unbalanced, which risked biasing the training by putting more weight on the abundant cell types during error minimisation. In this strategy, we generated new training pairs by randomly dropping 50% of the cells from the rabbit neighbourhoods multiple times, while leaving the paired mouse neighbourhoods unchanged. The amount of augmentation was set independently for each cell type, and chosen such that the number of neighbourhoods coming from each cell type was roughly equal after augmentation. Following this rule, we performed enough augmentations such that the number of examples from the most common cell type (erythrocytes) was multiplied by 2.

We also limited ourselves to a panel of 200 one-to-one orthologs for the prediction task. The panel was obtained from the mouse atlas using the function *gene_search* from the *geneBasisR* R package^57^. Limiting the number of genes helped to decrease computation time and memory requirements while retaining important biological information. We checked the latter by inspecting the “cell neighbourhood score” (Fig. S6c) of the package which estimates the distortion of cell-cell distances between the original dataset with all genes, and the dataset subsetted to the gene panel. As this metric is very costly to run on the entire dataset, we calculated it on a subset of the mouse atlas which contained 10% of the cells from each cell type.

Predictions from the random forest were evaluated using several metrics, alongside baseline predictions where mouse neighbourhoods were approximated by the expression profiles of their matched rabbit neighbourhoods. The coefficient of determination (R²) served as the primary goodness-of-fit metric, computed using *scikit-learn*’s *r2_score* function. Gene variability ranking was derived from the gene variance returned by the *model_gene_variance* function in *scranpy*^58^ (v0.1.3), applying the default LOWESS smoothing span of 0.3.

We integrated the neighbourhoods predicted by the random forest with the mouse cells using *scranpy*’s *fastMNN* implementation. Integration proceeded from the oldest to youngest embryos, with the largest samples processed first for embryos of the same age. All predicted neighbourhoods were treated as a single sample and integrated last. Prior to integration, the data underwent multi-batch PCA, with each sample blocked independently to generate 50 principal components, which were then used for fastMNN with 20 neighbours for MNN search. The corrected PCs were visualised using the *UMAP* function from the *umap* Python package, using the parameters: minimum distance 0.7 and 20 neighbours.

## Code availability

RIMA is implemented as an open source R package available at: https://github.com/ma-jacques/RIMA.

## Supplementary Figures

**Figure S1:**
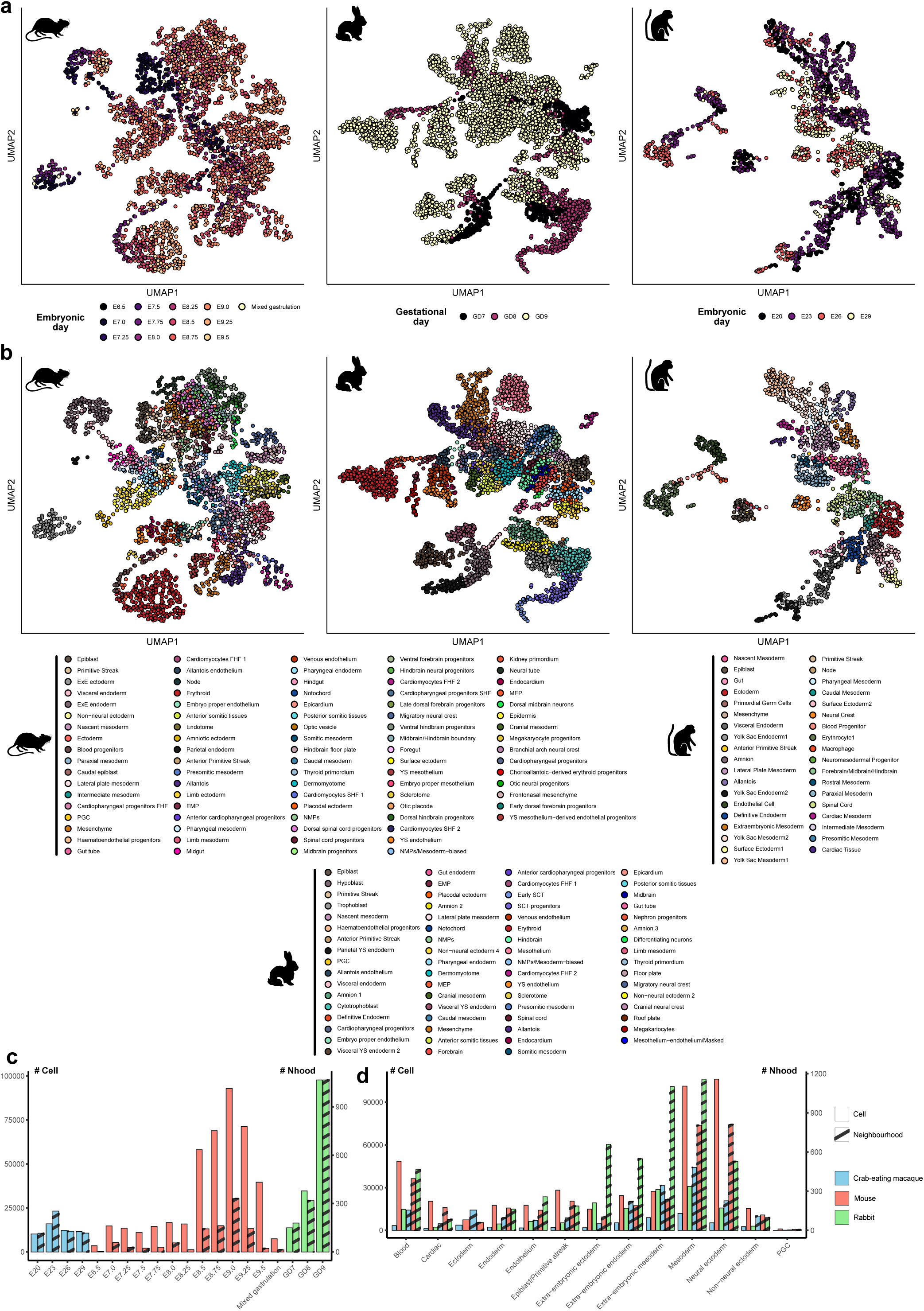
Presentation of the gastrulation atlases. **a-b**, UMAP visualization of cell neighbourhoods integrated with MNN for each species independently. Coloured according to cell type, and embryo stage neighbourhood annotations. **c-d**, Number of cells and neighbourhoods annotated per cell type (high-level systems, harmonized across species) and embryo stage.

**Figure S2:**
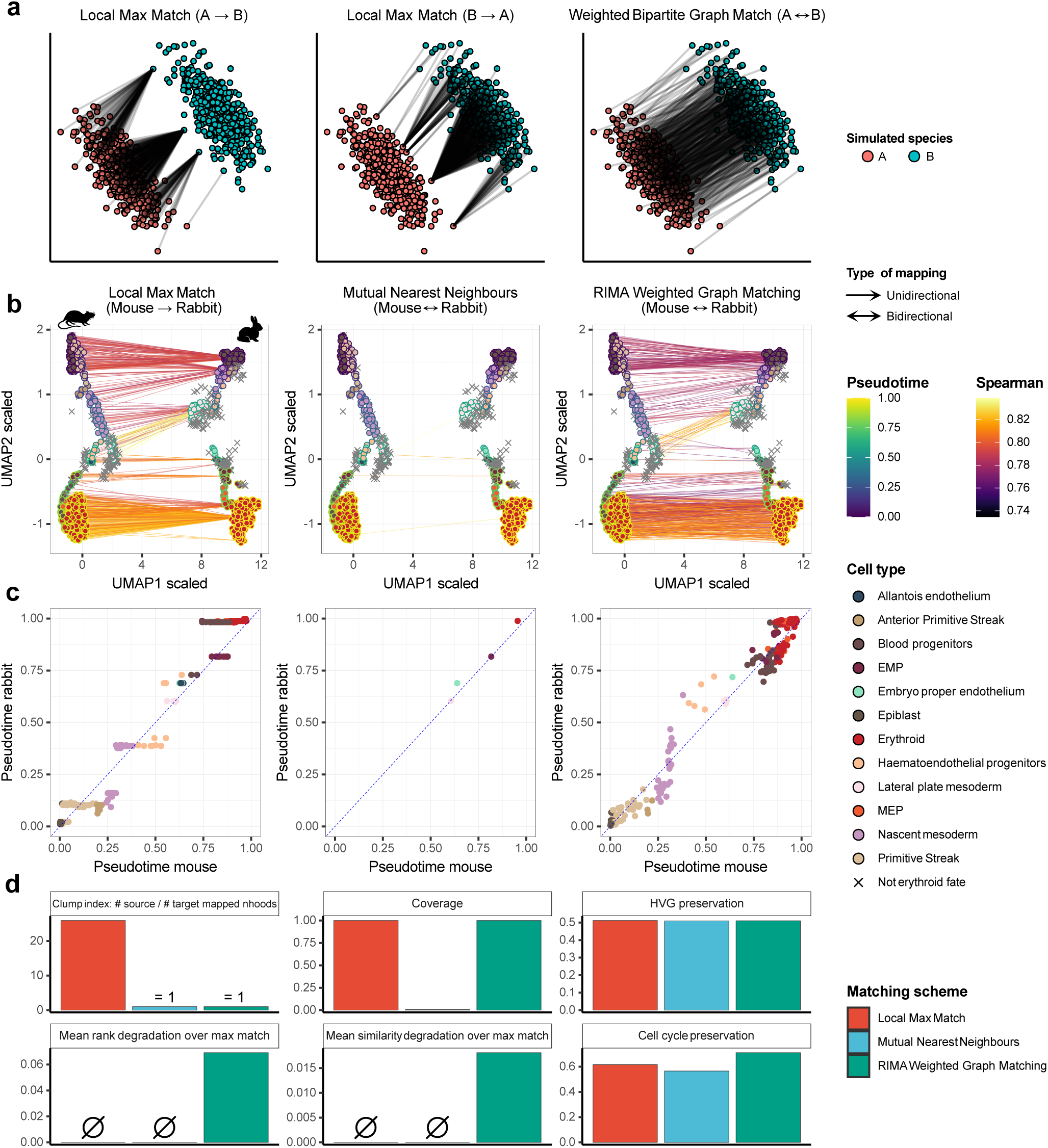
RIMA allows for accurate matching of cross-species biological states without integration. **a**, Comparison of 3 neighbourhood-neighbourhood matching methods for a simulated scenario where 2 matching clusters are shifted in expression space, as can be expected for unintegrated cross-species data. Left: Each red neighbourhood is matched to its closest neighbour (smallest L2 distance) in the blue cluster; Middle: same as left, but match blue on red instead; Right: Weighted Bipartite Graph Matching, same as used in RIMA. **b**, Comparison of matching schemes for the erythroid development trajectory in the mouse (left trajectory) and in the rabbit (right trajectory). The middle panel shows the pairs of neighbourhoods which are reciprocally maximally similar in both matching directions. **c**, Pseudotimes of the pairs of neighbourhoods defined in (**b**). **d**, Metrics associated with the matching schemes in (**b**). The clump index is the number of unique mouse neighbourhoods (source) over the number of rabbit neighbourhoods (target). Coverage indicates the number of neighbourhood pairs over the theoretical maximum number of 1:1 pairs; i.e. the number of neighbourhood pairs over the number of neighbourhoods in the smallest set (here, the rabbit set with n=675 neighbourhoods). The HVG preservation indicates the proportion of the top 2000 HVGs intersecting between the neighbourhoods matched in the rabbit and the ones matched in the mouse. The cell cycle preservation indicates the agreement of cell cycle annotations between the matched neighbourhoods; cell cycle phases were estimated with the cyclone function from the scran R package with the built-in mouse markers. Similarity degradation quantifies the difference in Spearman correlation between the paired rabbit neighbourhood and the mouse neighbourhood with which it is maximally correlated. Rank degradation quantifies the Spearman quantile rank of the paired rabbit neighbourhood and the mouse neighbourhood, where rank 1 is the maximally correlated pair and rank n (n=number of rabbit neighbourhoods) is the most dissimilar pair.

**Figure S3:**
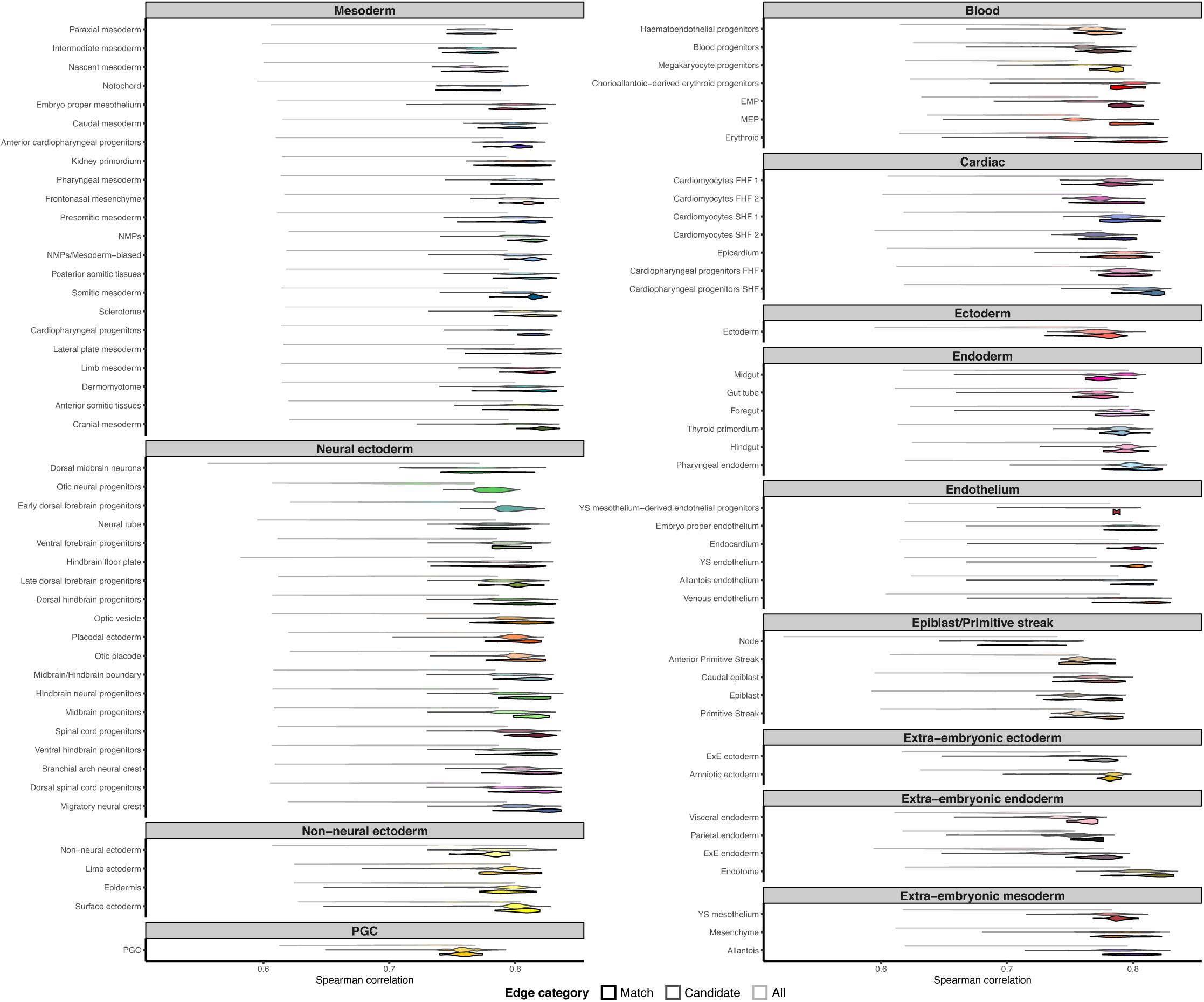
Distribution of similarities across cell types for different types of neighbourhood-neighbourhood edges: all, candidate matches, and actual matches. Candidate matches are the neighbourhood-neighbourhood edges that were deemed significant by RIMA, while actual matches regroup the subset of edges selected by RIMA in the final 1-to-1 matching.

**Figure S4:**
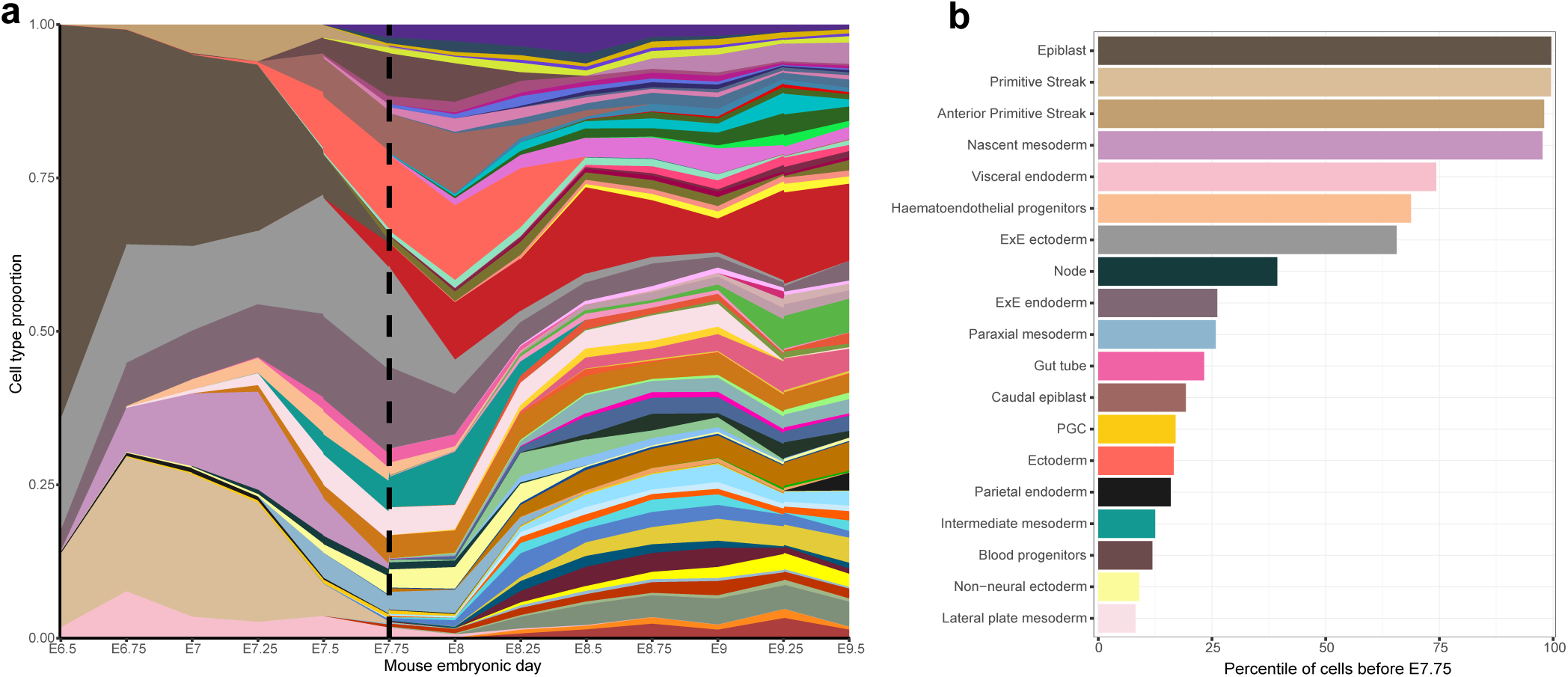
The gastrulation similarity bottleneck is characterized by the replacement of early undifferentiated tissues with a multitude of differentiated tissues. **a,** Proportion of cells at each mouse developmental stage. The vertical dashed line indicates the timing of mouse-rabbit similarity bottleneck. **b**, Mouse cell types for which at least 5% of the cells were observed before E7.75. Same colors as in Fig. S1.

**Figure S5:**
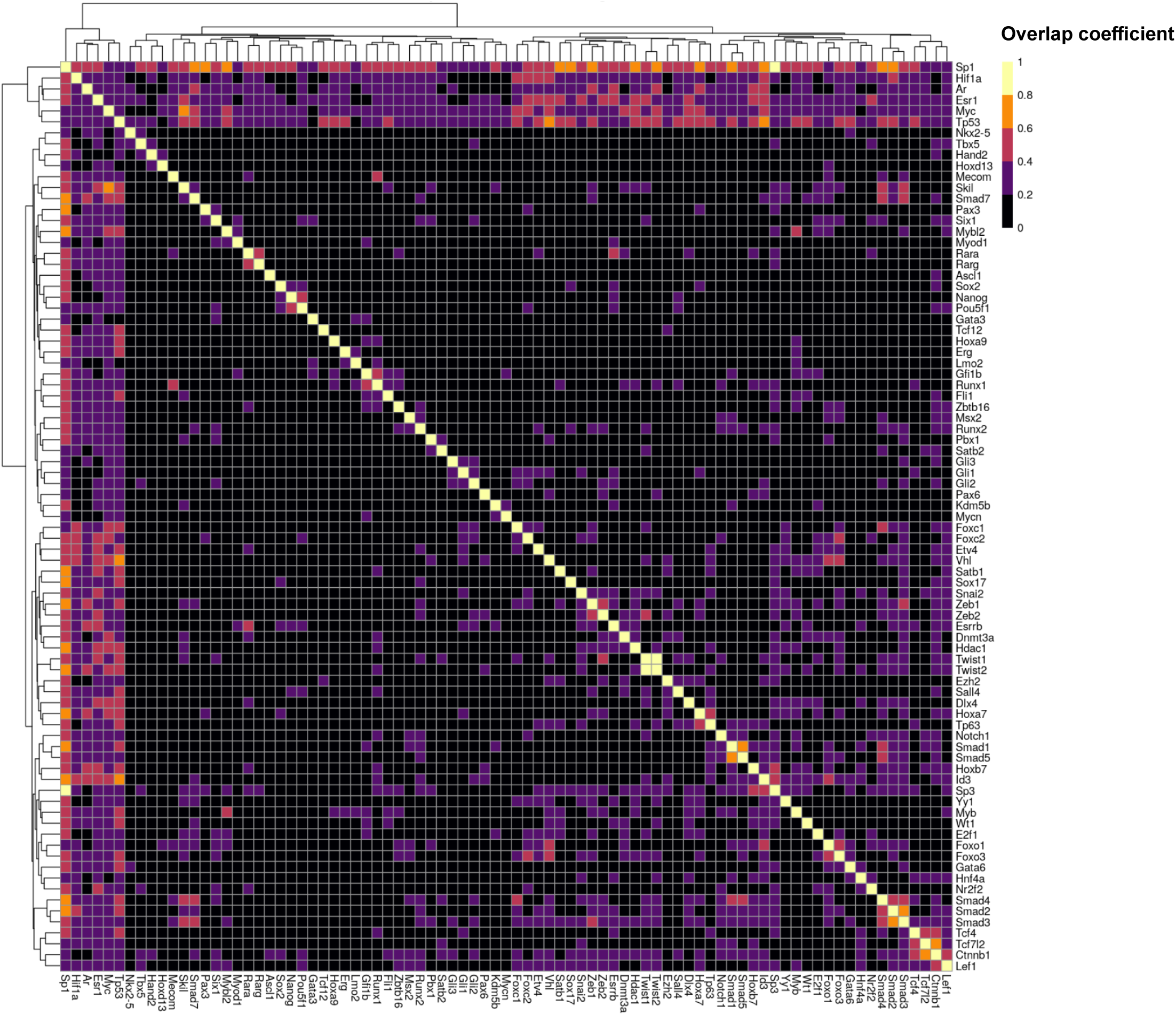
Core transcription factors’ regulons cover a large spectrum of target genes with little overlap. Heatmap of the overlap coefficients between the target genes of the Collectri regulons under the control of core TFs. The overlap coefficient between two regulons is defined as the number of target genes at their intersection, divided by the number of target genes in the smaller regulon.

**Figure S6:**
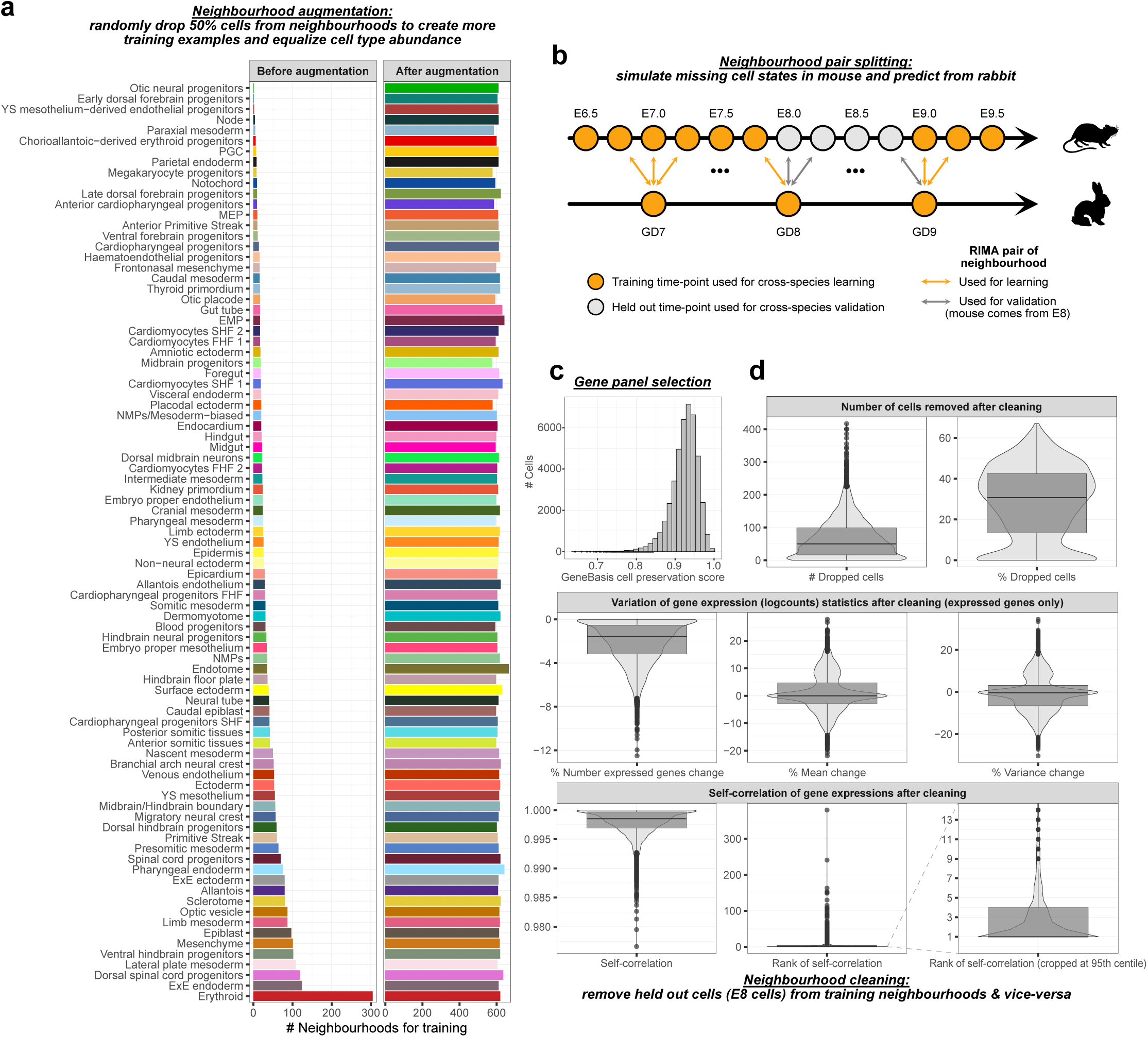
Set up for cross-species tuning learning and validation. **a**, Multiplication of training examples and correction for cell type imbalance through random drops of cells from training neighbourhoods. **b**, Schematic illustrating the formation of training and validation sets for cross-species prediction. RIMA’s neighbourhood matching was run on the complete mouse and rabbit datasets; then pairs where the mouse neighbourhood was annotated as coming from E8, E8.25, E8.5, or E8.75 were assigned to the validation set. **c**, Cell neighbourhood preservation metric of GeneBasis, calculated with the panel of 200 genes used for prediction.^57^ **d,** Metrics indicating: i) the effect of removing E8 mouse cells from mouse neighbourhoods assigned to training, and ii) the effect of removing non-E8 mouse cells from mouse neighbourhoods assigned to validation. These metrics were aggregated across i) and ii). Augmented neighbourhoods were not used to calculate the metrics. Self-correlation indicates the Pearson correlation of a neighbourhood average gene expression before and after cleaning. Correlations were also run between the cleaned neighbourhood and all the original mouse neighbourhoods to calculate self-correlation ranks. A rank of 1 means that the cleaned neighbourhood retains the original, non-cleaned neighbourhood as closest neighbour.

## Supplementary Tables

**Supplementary Table 1: Erythrocyte trajectories comparison, gene set enrichment analysis and CoPE scores**

**Supplementary Table 2: Regulon classification across mouse, rabbit and macaque according to shared conservation status**

**Supplementary Table 3: Per-gene statistics of cross-species prediction**

## Supplementary Note

### Supplementary Note 1: Matching single-cell profiles without cross-species integration

Single-cell transcriptomics measurements arise from a combination of biological and technical signals. While the latter is generally of less interest to biologists, it can represent a major source of variability that is difficult to disentangle from the biological signal of interest. The ability to cleanly separate technical from biological variation is worsened in the cross-species scenario, which introduces another source of major (biological) variation that is deeply entangled with technical effects as species are typically processed in separate sequencing batches. The lack of orthogonality between technical and biological signals is a major challenge for classical integration techniques, and aligning matching cell states across species can lead to a substantial and uncontrolled loss of biological signal.

In RIMA, we adopted a different approach to identify matching cell states across species while retaining as much biological signal as possible. We first defined biological states (neighbourhoods) on a per-species basis, allowing us to use classical integration techniques to safely remove batch effects across replicates (within a species). These states were then matched to close neighbours across species based on their gene expression counts, without additional correction.

A key feature of RIMA is that it matches entire sets of neighbourhoods at once, rather than individually pairing each neighbourhood with its closest counterpart. This feature is essential to correct for a strong artefact that appears when using nearest neighbours matching across unintegrated datasets. We illustrate this artefact in a simple simulation (Fig. S2a) where two matching, but shifted, clusters of points are mapped with the nearest neighbour approach. This approach leads to a heavily clumped matching, where only the points on the nearest periphery of each cluster are paired.

Although this coarse matching may still capture important biological variations (e.g. it still matches the right cell types; see Fig. S2b), it fails to account for other, more subtle, axes of variation within the clusters. This is visible when comparing the trajectories of erythrocyte development between mouse and rabbit (Fig. S2b-d). In this matching, the pseudotime of the matched neighbourhoods is correlated but is represented as a strong step-pattern, indicative of a loss of local variation/signal. This loss obscures subtle cross-species differences and would not allow the successful reconstruction of gene expression dynamics, such as the transcriptional boost of MURK genes (Fig. 3f).

## Acknowledgements

We would like to thank A. Yang, D. Kunz and V. Lorenzi for their feedback during the method development; and M. Barile for our discussions about the MURK genes dynamics development.

We would like to thank the European Molecular Biology Organization for supporting this project through a postdoctoral fellowship (ALTF 459-2022), as well as the Wellcome foundation for their support through the grants 220379/B/20/Z, 226795/Z/22/Z and 309075/Z/24/Z. J.C.M. acknowledges core funding from Cancer Research UK (C9545/A29580) and the European Molecular Biology Laboratory.

For the purpose of open access, the authors have applied a CC-BY public copyright license to any Author Accepted Manuscript version arising from this submission.

